# A lipid-mTORC1 nutrient sensing pathway regulates animal development by peroxisome-derived hormones

**DOI:** 10.1101/2021.05.15.444169

**Authors:** Na Li, Beilei Hua, Qing Chen, Meiyu Ruan, Mengnan Zhu, Huali Shen, Li Zhang, Huanhu Zhu

## Abstract

Animals have developed many signaling mechanisms that alter cellular and developmental programs in response to changes in nutrients and their derived metabolites, many of which remain to be understood. We recently uncovered that glucosylceramides, a core sphingolipid, act as a critical nutrient signal for overall amino-acid level to promote development by activating the intestinal mTORC1 pathway. However, how the intestinal GlcCer-mTORC1 activity regulates development throughout the whole body is unknown. Through a large-scale genetic screen, we found that the peroxisomes are critical for antagonizing the GlcCer-mTORC1-mediated nutrient signal. Mechanistically, deficiency of glucosylceramide, inactivation of the downstream mTORC1 activity, or prolonged starvation relocated peroxisomes closer to the intestinal apical region to release peroxisomal-beta-oxidation derived hormones that targeting chemosensory neurons to arrest the animal development. Our data illustrated a new gut-brain axis for orchestrating nutrient-sensing dependent development in *Caenorhabditis elegans*, which may also explain why glucosylceramide and peroxisome become essential in metazoans.

## Introduction

Animals have evolved sophisticated regulatory systems that sense the availability of environmental nutrients and modify their developmental, metabolic and behavioral programs accordingly. Extensive work in the last two decades has established mTORC1 as a central hub for sensing the level of a number of nutrients including amino acids (hereafter abbreviated as AA) in yeast and metazoans(Gonzalez and Hall, 2017; Kim and Guan, 2019; Wolfson and Sabatini, 2017). Genetic analyses using animal models have generated phenotypes that are consistent with mTORC1 being a critical nutrient-sensing system. Specifically, knockout of the major mTORC1 components, mTOR or RAPTOR, in mice caused early embryonic lethality shortly after implantation, when embryos must utilize environmental nutrients for development (Guertin et al., 2006). Such a phenotype mimics the temporary fasting-induced embryonic lethality (Gebhardt, 1969). Studies in invertebrate organisms such as *Drosophila melanogaster* or *C. elegans* also showed that deficiency of mTOR or RAPTOR orthologs caused developmental arrest and led to death at the postembryonic stage, which phenotypically mimicked the complete or AA starvation-induced development lethality(Jia et al., 2004; Long et al., 2002; Oldham et al., 2000; Qi et al., 2017). Extensive research in the field has also generated abundant mechanistic data to describe how mTORC1 senses specific dietary nutrients, including the roles of SLC38A9, Sestrin or CASTOR in sensing specific amino acids through physical interactions and consequently activates mTORC1(Chantranupong et al., 2016; Gu et al., 2017; Peng et al., 2014; Rebsamen et al., 2015; Tan et al., 2017). However, much of the mechanistic findings were made in cultured mammalian cells and have not been demonstrated to play critical roles in animal models under physiological conditions. For example, overexpression of conserved AA sensors such as Sestrin, which was expected to block the mTORC1 activity, did not cause similar early developmental lethality in flies, worms or mice (Kim et al., 2020; Peng et al., 2014; Yang et al., 2013). In addition, other reported AA sensors, such as SLC38A9 or CASTORs, are missing in invertebrate genomes(Saxton and Sabatini, 2017; Wolfson et al., 2016). Therefore, how mTORC1 senses AA in animals under specific physiological conditions (such as fasting) remains unclear. For multiple reasons, effective *in vivo* assay systems are not only valuable for verification of models raised by studies in cultured cells but perhaps also critical in discovering unknown mechanisms. For example, in multi-cellular animals, regulation in response to changed nutrient availability commonly involves sensing specific nutrients in specific tissues (such as intestinal cells or sensory neurons) as well as cross-tissue communications under physiological conditions. These important features are missing in cultured mammalian cells. In addition, commonly used immortalized cells are expected to be severely compromised in nutrient sensing capacity, as many well-known tumor suppressors and oncogenes including genes in the mTORC1 pathway play critical roles in major nutrient response signaling/regulatory pathways(Cui et al., 2013; Dazert and Hall, 2011; Saxton and Sabatini, 2017).

We and others previously found that monomethyl branched-chain fatty acid (mmBCFA), and its endogenous sphingolipid metabolite d17isoGlcCer, are critical for *C. elegans* postembryonic development (Kniazeva, Crawford et al. 2004, Kniazeva, Euler et al. 2008) (Zhu, Shen et al. 2013, Kniazeva, Zhu et al. 2015, Zhu, Sewell et al. 2015). Interestingly, deficiency of these lipids in whole animals, or even only in the intestine, will arrest development at the early postembryonic stage, which phenotypically copied the developmental arrest under complete starvation(Marza et al., 2009; Zhu et al., 2013). Moreover, activation of the conserved mTORC1 pathway by mutating its negative regulator *nprl-3* (orthologue of mammalian NPRL3) or constitutively activating key mTORC1 components restored the development of d17isoGlcCer-deficient animals (Zhu, Shen et al. 2013). Furthermore, we recently found that mmBCFA-GlcCer critically mediated mTORC1 sensing of the overall AA abundance during the *C. elegans* postembryonic development (Zhu et al., 2020). In addition, in mammalian tissue cultured cells, AA or mmBCFA supplementation could activate mTORC1 in a GlcCer dependent manner(Zhu et al., 2020). Finally, similar to mTORC1 mutants, embryos of UGCG (GlcCer synthase) knockout mice also grew smaller and eventually die shortly after implantation(Yamashita et al., 1999). These data strongly suggest that mmBCFA and its endogenous sphingolipid metabolite GlcCer critically mediate the mTORC1 dependent AA sensing during the early development in animals. However, since these findings would present a significant conceptual advance in AA/nutrient sensing by mTORC1, the new paradigm may not be firmly established without a better understanding of the mechanism downstream of this mmBCFA-GlcCer-mTORC1 signaling that is initiated in the intestine and regulates the global developmental program By a forward genetics screen in GlcCer-deficient *C. elegans* mutant, we surprisingly found that mutated peroxisomal proteins (peroxin) could significantly suppress the developmental defect and lethality of GlcCer and mTORC1 deficient animals. Our further analyses uncovered a previously unknown role of peroxisome in hormone production, secretion and developmental regulation orchestrated by the GlcCer/mTORC1-dependent food-sensing machinery.

## Result

### *prx-11* deficiency renders GlcCer dispensable in *C. elegans* development

To discover the specific pathway that mediates the nutrient-dependent developmental control downstream of GlcCer/mTORC1 in the nematode *C. elegans*, we carried out a genetic suppressor screen to identify mutations that restore the development of GlcCer-deficient animals. In *C. elegans*, there are three homologues of GlcCer synthase named *cgt-1, cgt-2* and *cgt-3* (Figure 1A). The *cgt-1/2/3* triple null mutant arrests development at the early first instar larval stage (L1 stage) (Figure 1A, B, E)(Marza et al., 2009; Zhu et al., 2013). Using the chemical ethyl methanesulfonate (EMS) to mutagenize *cgt-1/2/3* mutant animals (Figure 1B, Figure S1A), we identified 49 suppressor mutations that suppressed the growth arrest phenotype. A Q83stop mutation in the *prx-11* gene (determined by linkage analysis, Method) was among mutations that displayed the strongest suppression, as about 50% of *cgt-1/2/3; prx-11(sht1)* animals grew to adult and continuously propagated for infinite generations, even though the survivors grew slower and were smaller in adult size (Figure 1C-G). We determined that this suppressor caused a nonsense mutation (CAA-TAA, Glu83-Stop) (Figure 1H, Figure S1B, C, D, E). We then knocked down *prx-11* by RNAi in the *cgt-1/2/3; ex[WT cgt-1]* animals and found about 5% *cgt-1/2/3(−)* animals (n= 199) grew to adult, suggesting the *prx-11(sht1)* is casual to the suppression. The low suppression efficiency by *prx-11(RNAi)* was likely due to poor growth of *cgt-1/2/3(−)* animals on the RNAi carrier *E. coli* (HT115) strain (see below) and/or the partial reduction of gene activity by the RNAi treatment. To further confirm this, we generated two independent *prx-11* mutants [prx-11(CSR1) and prx-11(CSR2)] by CRISPR technology (Figure 1H)(see method). Each strain has a large deletion in the sole exon of *prx-11* that results in a frameshift (Figure 1H). Both mutations could effectively suppress the L1 arrest caused by *cgt-1/2(−);cgt-3(RNAi)* (Figure 1I). In this *cgt-3(RNAi)* treatment, an OP50-derived *E. coli* strain was used as a carrier to avoid the problems associated with the HT115 strain (Method) (Xiao et al., 2015). In addition, both the *cgt-1/2/3(−); prx-11(CSR1)* and *cgt-1/2/3(−); prx-11(CSR2)* quadruple mutants could complete their development (Figure 1J). Moreover, ectopic expression of the WT *prx-11* gene driven by its own promoter almost completely reversed the suppression effect of *prx-11(CSR2)* in the *cgt-1/2(−) cgt-3(RNAi)* background (Figure 1K). Therefore, we conclude that *prx-11* loss-of-function mutants [collectively referred to hereafter as *prx-11(-)*] could bypass the GlcCer deficiency-induced L1 arrest. Because these two CRISPR generated *prx-11* mutant strains were phenotypically healthier than the EMS-induced strain that likely contained many more background mutations, we mainly used them in the following studies.

**Figure 1.**
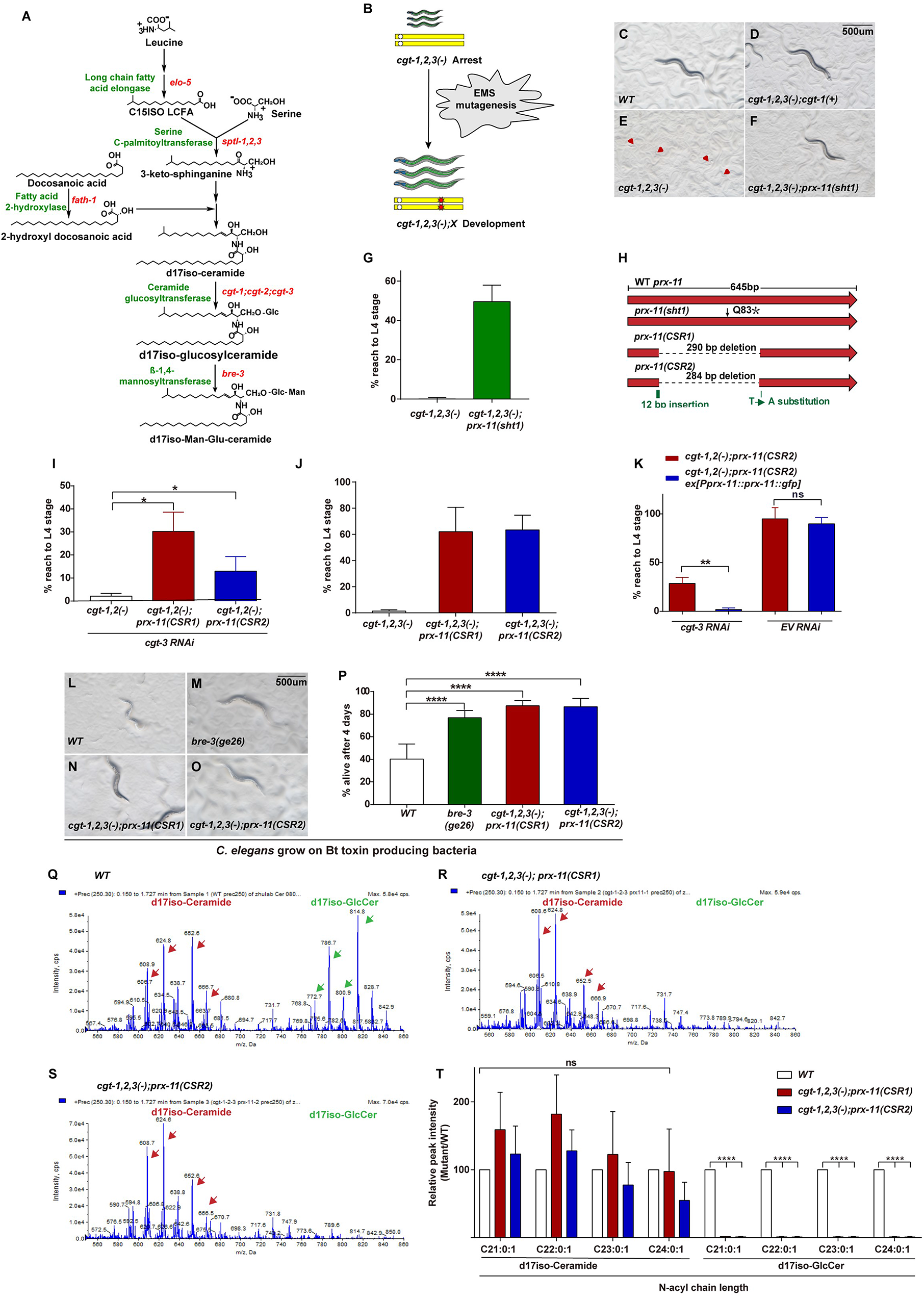
*prx-11* mutants restore the development of GlcCer-deficient animals without recovering the levels of GlcCer. (A). Sphingolipid biogenesis pathway in C. elegans, including the catalytic enzymes (green) and corresponding genes (red) used in this study. (B) The scheme of genetic suppressor screen for mutants that suppress GlcCer deficiency induced L1 arrest. (C-F) Images showing *cgt-1/2/3(−)* arrested at the L1 stage while *cgt-1/2/3(−)* carrying *WT cgt-1* or *prx-11* mutation reached adulthood; the quantitative data are presented in (G). (H) A cartoon illustration of the predicted structure and position of the *sht1* mutation in the *prx-11* gene. The molecular lesions of two *prx-11* alleles generated by CRISPR-CAS9 are also shown. (I) Two *prx-11* CRISPR mutants suppressed the L1 arrest phenotype of *cgt-1/2(−); cgt-3(RNAi)* and the L1 arrest phenotype of *cgt-1/2/3(−)* (J). (K) overexpression of WT *prx-11* under its native promotor reversed the suppression effect of *prx-11(−)*. (L-P) Image and quantitative data showing animals growing on the Bt-toxin Cry-5B producing bacteria BL21. Most WT animals (L) died in four days, but disruption of glucosylsphingolipid biosynthesis by *bre-3 (M), cgt-1/2/3(−) prx-11(CSR1)* (N) or *cgt-1/2/3(−) prx-11(CSR2)* (O) permitted survival past day 4. (Q-T) Shotgun mass spectrometry by precursor scan m/z = +250.3 showing there were many ceramides (red), but no detectable GlcCer (green) in *cgt-1/2/3; prx-11(−)* mutants (R, S). (T) Bar graphs from Multiple Reaction Monitoring (MRM) by UPLC-Mass spectrometry showing there were many of ceramides but only trace amount of GlcCer in *cgt-1/2/3; prx-11(−)* mutants.

It is unlikely that the *prx-11* mutations suppress the *cgt-1/2/3(−)* caused L1 arrest via restoration of endogenous GlcCer biosynthesis because *cgt-1, 2* and *3* are the only known enzymes to produce GlcCer in *C. elegans* (Marza et al., 2009). To confirm this, we first tested whether *cgt-1/2/3(−); prx-11(−)* quadruple mutants are resistant to Bt (Bacillus Thuringiensis) toxin Cry-5B produced from *E. coli* BL21Marroquin et al. (2000). A previous work showed that mutations that block the glucosylsphingolipid biosynthetic pathway (e.g., *bre-3*; see Figure 1A) could suppress the toxicity of Cry-5B toxin in *C. elegans* (Griffitts et al., 2001). Indeed, we found that, on the Cry-5B producing bacteria, both *cgt-1/2/3(−); prx-11(−)* quadruple mutants survived as well as a *bre-3(null)* mutant (Figure 1L-P), consistent with the notion that glucosylsphingolipid biosynthesis was blocked in *cgt-1/2/3(−); prx-11(−)* quadruple mutants.

We then directly measured the GlcCer level using liquid chromatography–mass spectrometry (LC-MS) and found that GlcCer portions were almost complete missing in *cgt-1/2/3(−); prx-11(−)* quadruple mutants, while the ceramide portion (the substrate of CGT-1/2/3 for GlcCer biosynthesis) was still abundant (Figure 1Q-T, Figure S1F). These data indicate that *pxr-11(−)* restored the development of *cgt-1/2/3(−)* without recovering GlcCer biosynthesis. In other words, PRX-11 may act downstream of GlcCer to suppress the developmental process.

### Peroxisome dysfunction suppresses developmental arrest caused by GlcCer deficiency

We then investigated how *prx-11(−)* suppressed the developmental arrest of GlcCer-deficient animals. PRX-11 is the *C. elegans* homolog of the PEX11, a conserved peroxisomal protein from yeast to human. We overexpressed the *prx-11* gene with a C-terminal-fusion GFP driven by its own promoter and found PRX-11::GFP was expressed across the whole body (Figure 2B, E). Within the cell, PRX-11 showed a punctate expression pattern that resembled the typical peroxisomal localization. The peroxisomal localization was confirmed by co-expressing an mCherry-tagged PMP-2 (a homologue of mammalian PMP70)(Lee et al., 2014) (Figure 2A, D), a conserved peroxisomal membrane protein as the peroxisome marker, and observing the co-localization of PRX-11 with PMP-2 (Figure 2C, F).

**Figure 2.**
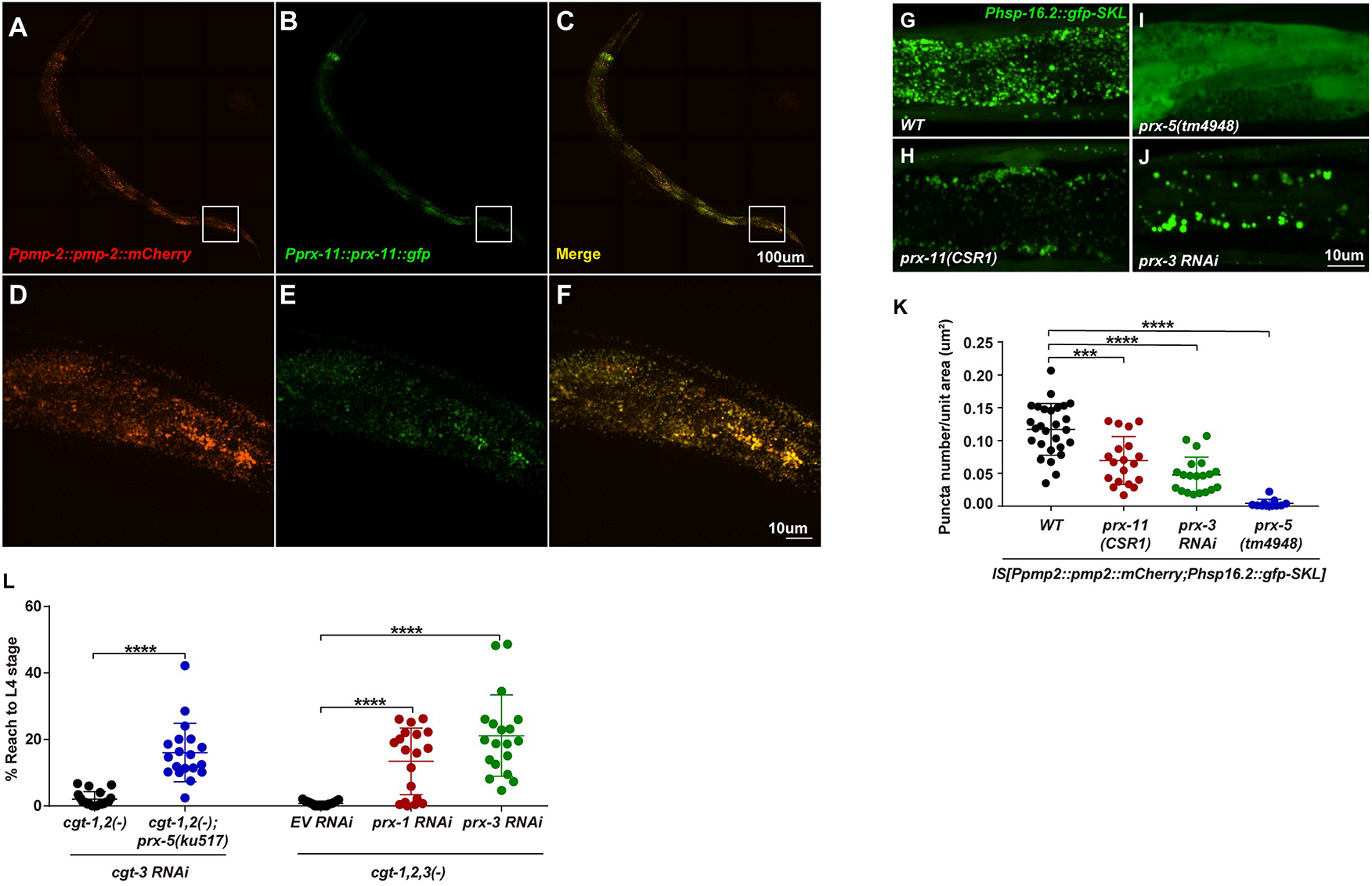
Peroxisomal dysfunction suppressed GlcCer deficiency induced developmental arrest. (A-F) Fluorescent microscopy images showing that PRX-11∷GFP (B) is ubiquitously expressed in *C. elegans* and colocalized with peroxisomal membrane protein PMP-2::mCherry (A). Enlarged images shown in D-F. (G-J) Fluorescent microscopy images showing the peroxisomes marked by peroxisomal targeting protein GFP-SKL. Compared to WT (G), the peroxisomal numbers in *prx-11(−)* (H) and *prx-3^RNAi^* (J) were dramatically decreased. In *prx-5(−)* (I), the GFP-SKL protein was no longer targeted to peroxisomes. The quantitative data are shown in K. (L) Bar graphs showing *prx-5(ku517)*, *prx-1^RNAi^* or *prx-3^RNAi^* also suppressed GlcCer deficiency induced L1 arrest.

The homologues of PRX-11 (PEX11) in budding yeast and mammals play important roles in peroxisomal division/proliferation (Li et al., 2002; Li and Gould, 2002). Using a previously reported peroxisomal target protein, GFP-SKL, to mark the peroxisome (Thieringer et al., 2003; Wang et al., 2013), we found that *prx-11(−)* severely reduced the number of peroxisomes, indicating PRX-11 is also involved in peroxisomal proliferation in *C. elegans* (Figure 2G, H, K). Therefore, we hypothesized that the suppression effect of *prx-11(−)* on *cgt-1/2/3(−)*-induced L1 arrest was due to reduced peroxisomal function, which could be tested by using other peroxisomal mutants. PRX-5 (ortholog of human PEX5) is a critical peroxisomal protein to transport enzymes into peroxisomal matrix(Thieringer et al., 2003; Wang et al., 2013). In *prx-5(−)*, GFP-SKL is not be transported into the peroxisome and thus causes peroxisomal dysfunction as previously reported (Figure 2I) (Thieringer et al., 2003; Wang et al., 2013). PRX-3 (ortholog of human PEX3) is another critical peroxisomal protein that is responsible for peroxisome *de novo* biogenesis(Yuan et al., 2016). With *prx-3(RNAi)*, the peroxisomal number was also dramatically reduced, and the remaining peroxisomes were abnormally large (Figure 2J, 2K). We found that a *prx-5(−)* mutation (Wang et al., 2013), or RNAi of either *prx-3* or *prx-1,* partially suppressed the *cgt-1/2/3(−)* induced L1 arrest (Figure 2L). These data indicate that deficiency in peroxisomal biogenesis or general function suppresses GlcCer deficiency-induced developmental arrest.

### Peroxisome acts downstream of sphingolipid to regulate development

We next asked whether peroxisome deficiency suppressed the GlcCer deficiency by altering the activity of the d17isoGlcCer biosynthetic pathway. We previously found that the *de novo* monomethyl branched-chain fatty acid (mmBCFA) or mmBCFA-related sphingolipid pathways play essential roles in early postembryonic development (Figure 1A). Deficiency in these pathways mimicked a starved environment and caused *C. elegans* to arrest at early L1 stage and display foraging defects (Kniazeva et al., 2004; Kniazeva et al., 2008; Kniazeva et al., 2015; Zhu and Han, 2014; Zhu et al., 2015; Zhu et al., 2013). Interestingly, we found *prx-11(−)* could also effectively suppress the developmental phenotypes that arise from RNAi knockdown of the *de novo* mmBCFA or sphingolipid biosynthesis pathway [*elo-5* (mmBCFA elongase), *fath-1* (fatty acid 2-hydroxylase) and *sptl-1* (serine-palmitoyl transferase)](Figure 3A). These data indicate *prx-11* acts downstream of GlcCer to regulate development.

**Figure 3.**
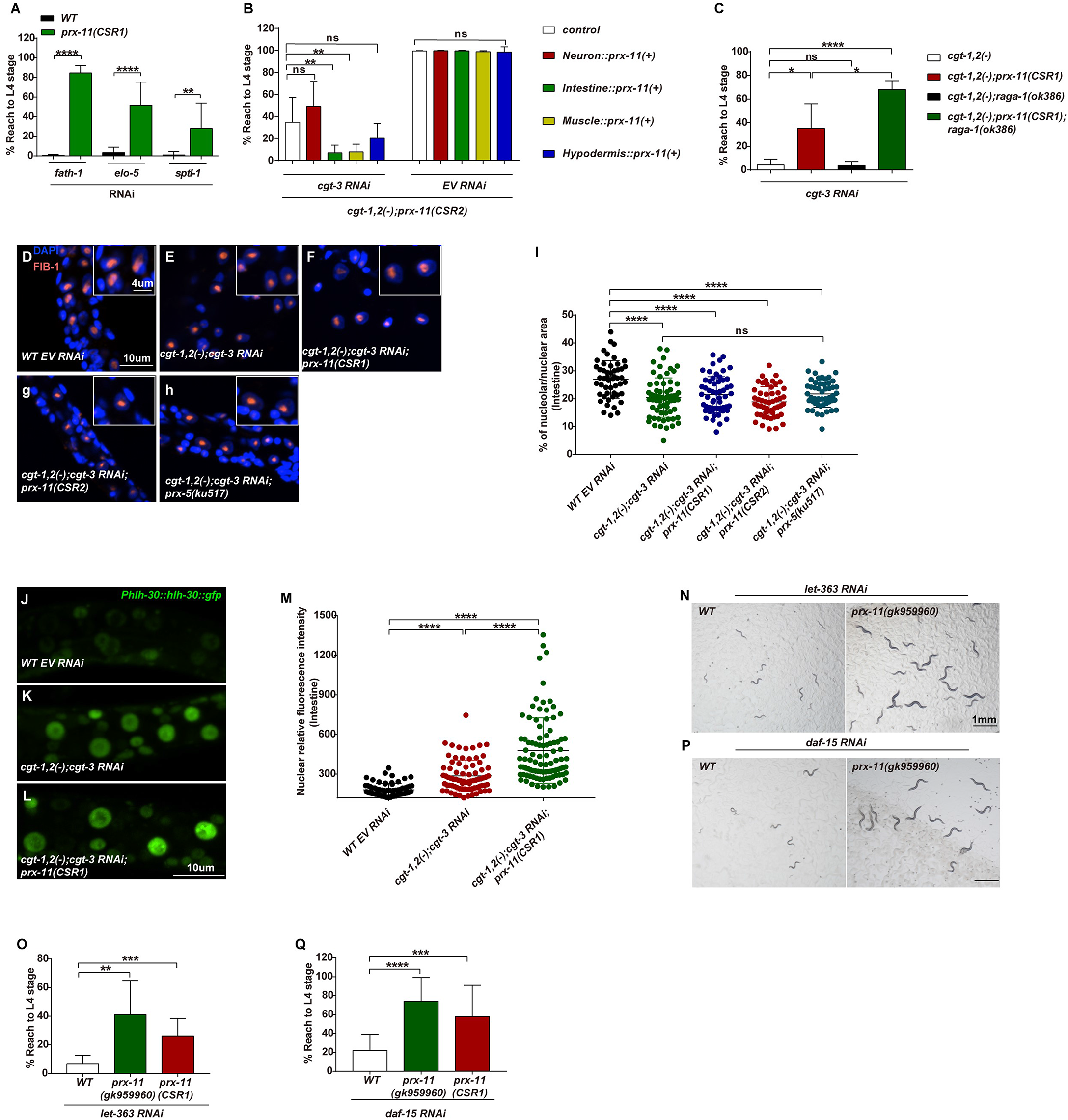
Peroxisomal mutants act downstream of GlcCer/mTORC1 pathway to promote *C. elegans* development. (A-B) Bar graphs showing percentage of animals that reached the L4 stage. (A) *prx-11(CSR1)* suppressed the developmental arrest caused by RNAi knockdown of various genes in the *de novo* sphingolipid pathway. (B) The development of *cgt-1/2/3(−) prx-11(−)* animals was suppressed by intestinal or muscle-specific restoration of *PRX-11*. (C) Bar graph showing that deficiency of mTORC1 by *raga-1* mutant did not reverse the developmental restoration of *prx-11(−)* in GlcCer-deficient animals. (D-H) Representative fluorescent microscopic images showing the relative nucleolar size (a marker of mTORC1 activity) of intestinal cells. It was significantly reduced in GlcCer-deficient arrested animals (E). Restoration of the development by *prx-11(−)* (F, G) or *prx-5(−)* (H) did not recover the nucleolar size. The statistical data are shown in (I). (J-M) Representative fluorescent microscopic images (J-L) and quantitative data (M) showing the nuclear intensity of HLH-30::GFP (a marker of reduced mTORC1 activity). GlcCer-deficient animals showed an increased nuclear HLH-30::GFP (K), while *prx-11(−)* did not suppress the nuclear HLH-30::GFP intensity in GlcCer deficient animals (L). (N-Q) Microscopy images (N, P) and bar graphs (O, Q) showing *prx-11(−)*— suppressed *let-363/mTOR* (N, O) or *daf-15/RAPTOR* (P, Q) RNAi induced developmental arrest, respectively.

Because the *de novo* mmBCFA and sphingolipid pathways mainly act in the intestine to promote *C. elegans* development (Kniazeva et al., 2008; Zhu et al., 2015; Zhu et al., 2013), we examined whether *prx-11* also acts in the intestine to execute its negative regulatory role. As expected, we found that restoration of WT PRX-11 expression in the intestine, but not in neurons or the hypodermis, almost completely reversed the suppression effect of *prx-11(−)*(Figure 3B). Interestingly, PRX-11 restoration in the muscle also showed the same suppression-reversion effect, suggesting *C. elegans* needs to keep peroxisomal function low in either or both muscle and intestine to restore development in GlcCer-deficient animals. These data suggest a cross-tissue interplay between the intestine and muscle to coordinate the developmental regulatory process (further addressed below)(Zhang et al., 2011; Zhu et al., 2015).

### The peroxisome acts downstream of mTORC1 to repress GlcCer-promoted development

We previously found that deficiency in *de novo* mmBCFA/GlcCer biosynthesis caused L1 arrest by inactivating the mTORC1 pathway (Blackwell et al., 2019; Entchev et al., 2008; Gonzalez and Hall, 2017; Kim and Guan, 2019; Wolfson and Sabatini, 2017; Zhu and Han, 2014; Zhu et al., 2015; Zhu et al., 2013). In addition, we found that mmBCFA and GlcCer critically mediated the sensing of overall amino acid (AA) abundance by mTORC1 under severe AA scarce condition in *C. elegans* and mammalian cells (Zhu et al., 2020). We thus examined the relationship between the peroxisomal function and the mTORC1 activity. We found that blocking mTORC1 by mutating *raga-1*(ortholog of mammalian RagA/B) did not affect the suppression effect of *prx-11(−)* (Figure 3C). Moreover, the reduced mTORC1 activity, as indicated by the decreased intestinal nucleolar size (Ma et al., 2016; Tiku et al., 2017; Wu et al., 2018) and by the increased nuclear intensity of *C. elegans* TFEB homolog HLH-30::GFP (Blackwell et al., 2019; O’Rourke and Ruvkun, 2013), in *cgt-1/2/3(−)* animals, was not recovered by the presence of *prx-11(−)* or *prx-5(−)* mutations (Figure 3D-M). In addition, we also found *prx-11(−)* could also significantly suppress larval arrest caused by RNAi of either *let-363/mTOR* or *daf-15/RAPTOR* (Figure 3N-Q). These data suggest that the peroxisome mainly acts downstream of mTORC1 to regulate *C. elegans* development.

### GlcCer/mTOR deficiency caused abnormal accumulation of peroxisome at the intestinal apical region

To understand how defects in peroxisomal biogenesis or function suppress L1 arrest induced by GlcCer deficiency, we first determined that neither GlcCer deficiency-induced, nor fasting-induced, developmental arrest was due to increased peroxisomal number or activity (Figure S4N, S4O). We further ruled out that the L1 arrest is caused by increased ROS stress (Schrader and Fahimi 2006), since neither mitochondrial UPR (measured by HSP-6::GFP) nor general ROS activity was elevated in GlcCer-deficient animals (Figure S4A-L). Moreover, addition of a ROS inhibitor (N-acetyl-l-cysteine) could not reverse the arrest (Figure S4M).

We then hypothesized that peroxisomes may play a specific regulatory role under GlcCer deficiency. Strikingly, we found that in *cgt-1/2/3(−)* animals, most intestinal peroxisomes, marked by a peroxisomal matrix protein DAF-22∷GFP(Joo et al., 2009), accumulated in the apical region (Figure 4A-B). The notion that GlcCer deficiency causes apical peroxisomal accumulation is also consistent with a reported observation that apical peroxisomal accumulation was observed with RNAi knockdown of *fath-1* (fatty acid 2-hydroxylase)(Li et al., 2018). FATH-1 is involved in biosynthesis of 2-hydroxyl very long chain fatty acid that is present in major GlcCers in *C. elegans* (Chitwood et al., 1995; Zhang et al., 2011; Zhu et al., 2013)(Figure 1A). Moreover, we found that apical peroxisomal accumulation was also observed under several other RNAi conditions that disrupt the GlcGcer-mTOR-mediated nutrient sensing pathway, including knockdown of genes in apical polarity machinery *chc-1* or *aps-1*, *let-363*/mTOR or *daf-15*/raptor (Zhang et al., 2011; Zhu et al., 2015), as well as under prolonged fasting (Figure 4A, 4B, 4D-J). The apical peroxisomal accumulation phenotype was prominently suppressed by the *prx-11(−)* mutation (Figure 4C, 4K). These data indicate that disruption of the GlcCer-mTORC1 pathway, that induces developmental arrest, also causes peroxisomal apical accumulation.

**Figure 4.**
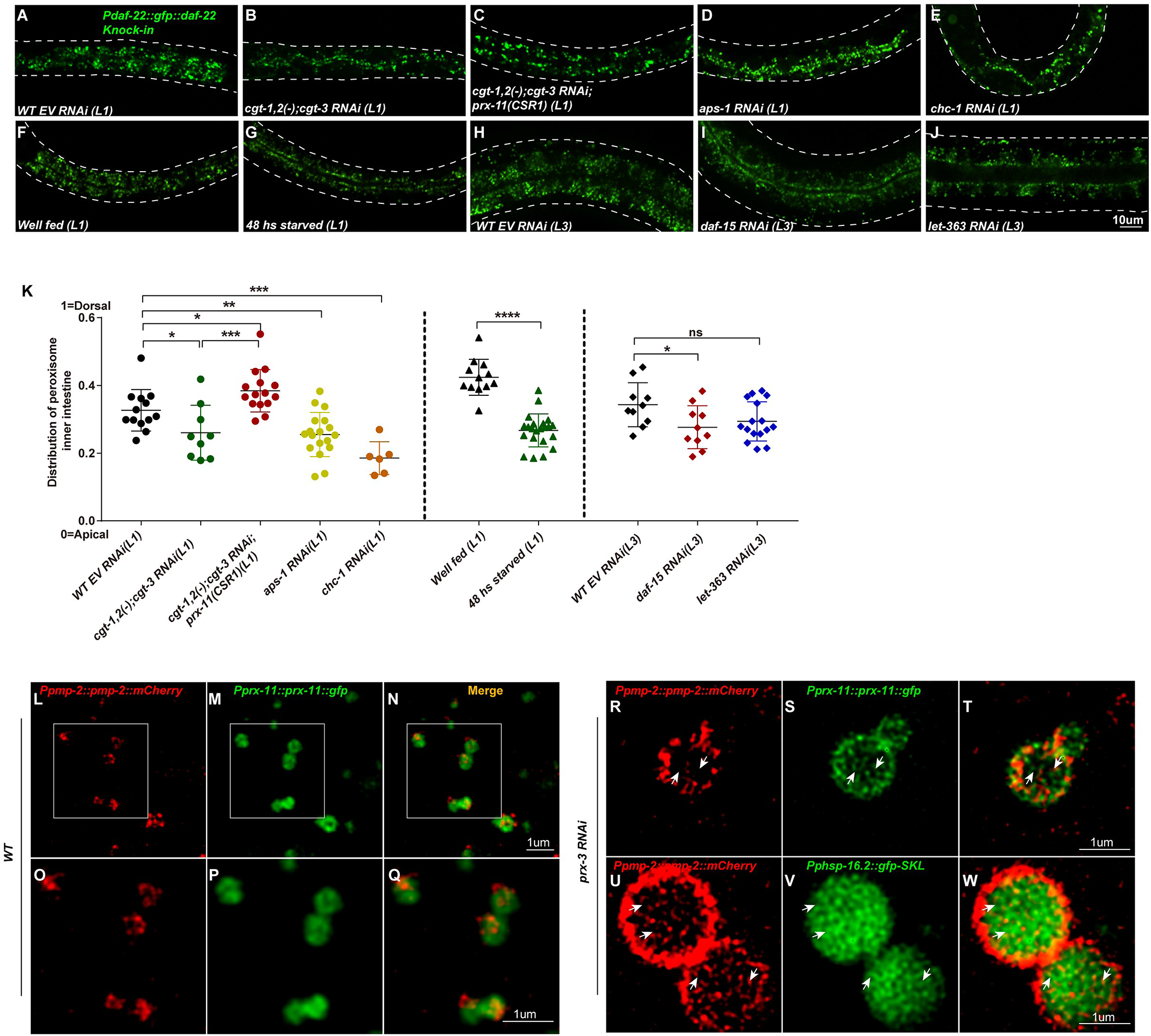
Peroxisomes accumulated in the apical region under GlcCer/mTORC1 deficiency or starvation. (A-K) Representative fluorescent images (A-J) and quantitative data (K) showing subcellular localization of peroxisomes marked by GFP∷SKL in C. elegans intestines. Compared to WT, disruption of GlcCer biosynthesis (*cgt-1/2/3^RNAi^*, [B]), vesicle trafficking pathway (by *aps-1^RNAi^* [D] or *chc-1^RNAi^* [E]), mTORC1 (*let-363^RNAi^* [I] or *daf-15^RNAi^* [J]) or complete starvation (G) caused the peroxisome to accumulate at the near apical region in the intestine. *prx-11(−)* restored the abnormal accumulation pattern in GlcCer-deficient animals (C). (L-Q) Super resolution fluorescent images showing PRX-11 (M, enlarged in P) co-localized with PMP-2 (L, enlarged in O) on the peroxisomal membrane. The merged image is shown in N (enlarged in Q). (R-W) Super resolution fluorescent images showing under *prx-3^RNAi^*, the PMP-2∷mCherry (R, U) and PRX-11∷GFP (S) also localized on the intralumenal vesicles, while peroxisomal matrix protein GFP-SKL formed many empty hollows (V). The merged images are shown in T, W.

### Peroxisomes contain intralumenal vesicles

A previous report suggested that PEX11 (yeast ortholog of PRX-11) plays an important role in extracellular secretion of small-molecule metabolites in filamentous fungi (Kiel et al., 2005). Therefore, based on our results described in Figure 4A-K, we hypothesized that these apical localized peroxisomes repressed developmental growth of *C. elegans* possibly by facilitating the secretion of certain metabolites that “signal” a poor-nutrient condition or metabolic status. Because we observed no peroxisomal matrix protein DAF-22 in the intestinal lumen in GlcCer deficient animals (Figure 4B), the direct and complete fusion of peroxisome with intestinal apical membrane to release such signal metabolites into the intestinal lumen seemed unlikely. Therefore, we hypothesized that there are intraperoxisomal vacuoles for the storage and release of signal metabolites, similar to synaptic vesicles in neurons(Lin and Scheller, 2000). Interestingly, a recent study reported that plant peroxisomes contain many intralumenal vacuoles (Wright and Bartel, 2020). However, there has been no reports of similar structures in animals, primarily due to much smaller sizes of animal peroxisomes (0.1-1 μm) that may not be readily detected by the standard confocal microscopy.

Therefore, we employed super-resolution stimulated emission depletion (STED) microscopy to explore such a possibility. Under STED, PRX-11::GFP and PMP-2 clearly marked the WT peroxisome membrane(Lee et al., 2014) (Figure 4L-Q), but the detailed intraluminal structure was not clear due to the small peroxisomal size (about 0.2-0.3μm). However, since we had found that *prx-3(RNAi)* could significantly enhance the peroxisome size (Figure 2J), we were able to make a surprising observation that PRX-11 and PMP-2 also localized in the peroxisomal lumen as puncta (Figure 4R-T, U, W). In addition, the peroxisomal matrix protein GFP-SKL was not evenly distributed in the peroxisomal lumen but displayed many intralumenal hollows (which lacked the GFP-SKL signal), suggesting the existence of intralumenal vesicles (Figure 4V, W). Taken together, these data suggest that there are peroxisomal intralumenal vesicles which may facilitate the storage and secretion of metabolites as a “poor nutrient hormone” under the regulation of GlcCer/mTORC1 pathway.

### Peroxisomal beta-oxidation derived ascarosides repress GlcCer/mTOR promoted development

We next searched for the potential peroxisomal-derived metabolites that might act as a “poor-nutrient hormone”. One of the major functions of the peroxisome is to execute beta oxidation, a process which generates many lipid metabolites. In *C. elegans*, several conserved proteins such as MOAC-1 (enoyl CoA hydratase), DHS-28 (3-hydroxyacyl CoA dehydrogenase), and DAF-22 (Thiolase) are critical enzymes responsible for peroxisomal beta oxidation. We found that RNAi knockdown of these genes all effectively suppressed the GlcCer deficiency-induced L1 arrest (Figure 5A), suggesting that peroxisomal beta oxidation is involved in generating the metabolites that act as a “poor-nutrient hormone”.

**Figure 5.**
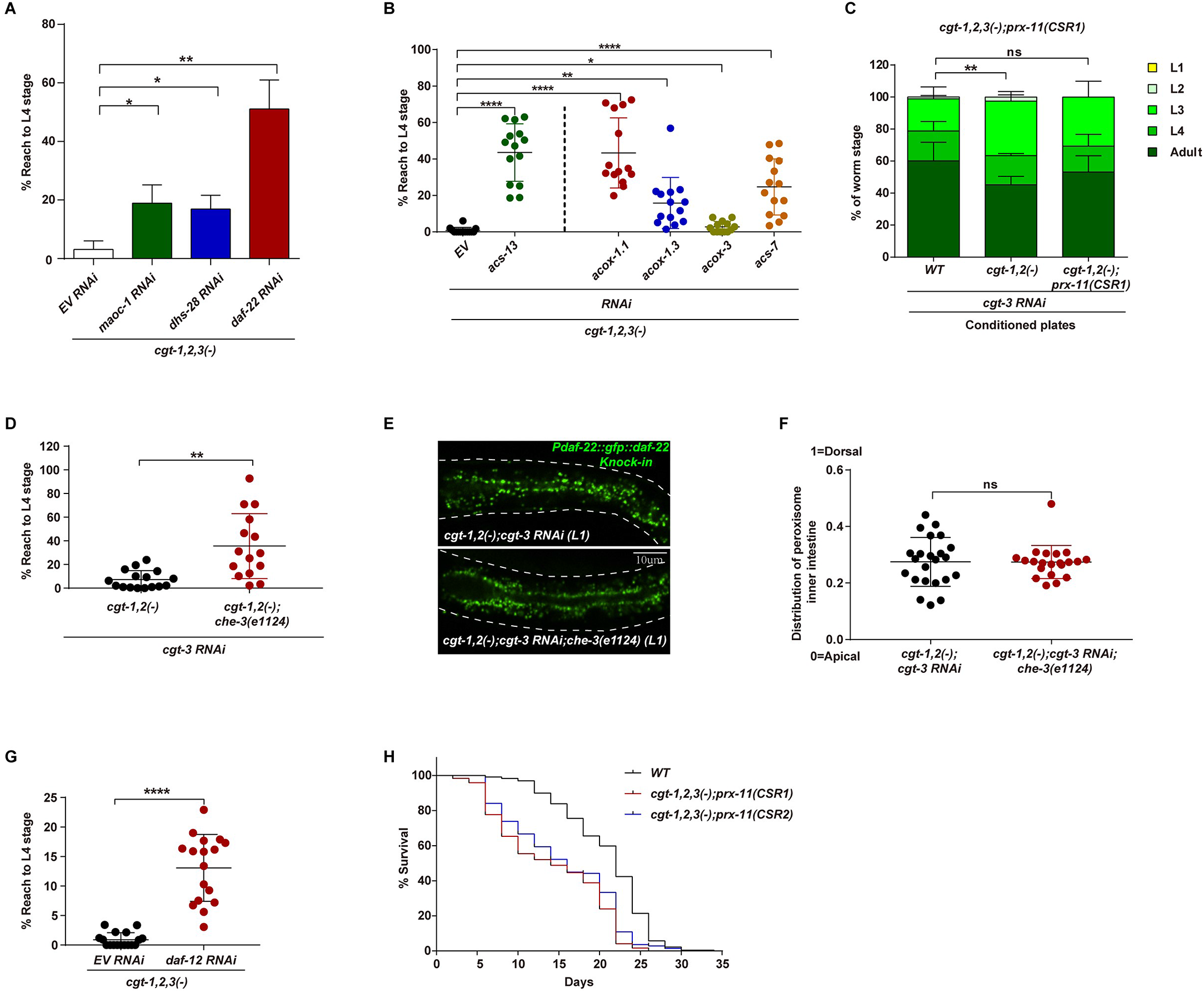
Peroxisomal beta-oxidation derived ascarosides repress GlcCer/mTOR promoted development. (A, B) Bar/dot graphs showing the percentage of GlcCer-deficient animals bypassed the developmental arrest. Disruption of peroxisomal β-oxidation genes *maoc-1*, *dhs-28 or daf-22* (A), medium chain acyl-CoA synthases *acs-13*, *acs-17* (B, left panel) suppressed GlcCer-deficient developmental arrest. Disruption of medium to short chain acyl-CoA oxidase *acox-1.1* or *acox-1.3*, or ocas#9/icas#9 biosynthetic gene *acs-7*, also suppressed GlcCer-deficient animals induced developmental arrest (B, right panel). (C) Bar graphs showing the medium secreted from *cgt-1/2/3* but not *cgt-1/2/3; prx-11(−)*, partially suppressed the development of *cgt-1/2/3; prx-11(−)* animals, indicating a GlcCer deficiency dependent pheromone was involved in the developmental regulation. (D) A dot graph showing the percentage of GlcCer-deficient animals bypassed the developmental arrest. *che-3(−)*, but not *osm-3(−)*, suppressed GlcCer-deficiency caused L1 arrest. (E, F) Representative fluorescent images (E) and quantitative data (F) showing subcellular localization of peroxisomes marked by GFP::SKL in *C. elegans* intestines. *che-3(−)* did not reverse the apical accumulation of peroxisomes in GlcCer-deficient animals. (G) A dot graph showing the percentage of GlcCer-deficient animals that bypassed the developmental arrest in *daf-12^RNAi^*. (H) Survival curves showing the percentage of L1 animals that survived under well-fed condition. WT animals survived better than *cgt-1/2/3(−)*; *prx-11(−) animals*.

In budding yeast *S. cerevisiae*, PEX11 was reported to be essential for the transportation of mid-chain fatty acids (MCFA, 8-12 carbon) from cytosol to peroxisome. These MCFAs were further catalyzed by MCFA acyl-CoA synthetase FAA2 before being β-oxidated to facilitate peroxisome proliferation(van Roermund et al., 2000). To determine whether PRX-11 dependent MCFA β-oxidation also played a vital role in the *C. elegans* developmental regulatory process, we knocked down *acs-13*, the closest homolog of yeast *faa2* by sequence alignment, and found it also significantly suppressed GlcCer induced developmental arrest (Figure 5B). These data suggest a short/medium chain FA or its metabolites, derived from MCFA β-oxidation, may act as the negative developmental regulatory hormone.

In *C. elegans*, one group of well-known secreted metabolites, which are peroxisomal-beta oxidation dependent, are ascarosides that consist of a fatty acid linked to a dideoxysugar ascarylose (von Reuss et al., 2012). They act as hormones/pheromones to regulate multiple developmental processes and behaviors of *C. elegans* including dauer entry and exit(Ludewig and Schroeder, 2013). A previous report showed that the level of short chain ascarosides were dramatically decreased in peroxisomal beta oxidation mutants such as MOAC-1, DHS-28, or DAF-22(von Reuss et al., 2012). Therefore, we hypothesized that the production of short chain ascarosides could be responsible for the GlcCer deficiency-induced developmental arrest. Several acyl-CoA oxidases (ACOXs) are known to initiate peroxisomal β-oxidation by dehydrogenating the acyl-CoA between the α and β carbon, and different ACOXs have different substrate specificities (Zhang et al., 2015). To understand which class of short chain ascarosides may contribute to the GlcCer-dependent developmental regulation, we first knocked down several acox genes in GlcCer-deficient animals. We found of *acox-1.1(F08A8.1)^RNAi^* restored the development at the level similar to that by DAF-22^RNAi^, while *acox-1.3(F08A8.3)^RNAi^* displayed a partial suppression (Figure 5B). RNAi of *acox-3(F58F9.7)^RNAi^* only had a minor suppression effect (Figure 5B). Because ACOX-1.1 deficient animals displayed dramatic decrease in C5, ∆C7 and ∆C9 ascarosides, while ACOX-1.3 deficient animals showed decrease ∆C7 ascaroside (Zhang et al., 2015) (Figure S5A), we reasoned that C5, ∆C7 and ∆C9 families could be the major developmental suppression ascarosides secreted in GlcCer-deficient animals. Interestingly, production of C5 ascarosides was reported to be significantly increased under fasting induced L1 arrest(Artyukhin et al., 2013), which is also consistent with our hypothesis. In addition to the core C5 ascaroside ascr#9 (Figure S5A), the C5 ascarosides family consists of several subtypes based on the additional moiety on the ascarylose, such as octopamine ascaroside osas#9 and indole ascaroside icas#9 (Figure S5A). The biosynthesis of these ascarosides completely depends on ACS-7(Artyukhin et al., 2013). Interestingly, we found acs-7^RNAi^ could also effectively restore the development of GlcCer-deficient animals (Figure 5B), even though the suppression effect was not as robust as that of *acox-1.1 ^RNAi^*. These data suggest osas#9 and icas#9 are also among those development-repressing metabolites.

We further tested the ability of ascarosides obtained from GlcCer-deficient animals in delaying the development in peroxisome defective [*prx-11(−)*] animals. We let *cgt-1/2(−) cgt-3^RNAi^* animals grow on plates for 2 days to secrete ascarosides first, then transferred fluorescent-labeled *cgt-1/2/3(−); prx-11(−)* L1 animals to the *cgt-1/2/3(−)* pre-conditioned plates (see method). Indeed, more *cgt-1/2/3(−); prx-11(−)* arrested at the early developmental stage *in cgt-1/2/3(−)* pre-conditioned plates (Figure 5C). The incomplete developmental arrest may be due to the low concentration of ascarosides in the pre-conditioned medium, or due to a role of these ascarosides as an endocrine hormone rather than a pheromone from the environment to regulate development (Ludewig et al., 2017). We then performed two more tests to distinguish between them. First, using LC-MS to identify related secreted ascarosides, we found C5 (ascr#9 and/or icas#9), ∆C7(ascr#7), ∆C9(ascr#3) and C9(ascr#10) ascarosides were both significantly upregulated in the excretomes (liquid culture supernatant extracts)(Figure S5 B, D, F, H, J, L, N) and worm body(Figure S5 C, E, G, I, K, M, O) of *cgt-1/2/3(−)* animals, and dramatically downregulated in the *cgt-1/2/3(−); prx-11(−)* animals (Figure S5P, 5Q), which suggests these ascarosides could be acting as pheromone or hormone to suppress *C. elegans* development. Second, we grew singled GlcCer-deficient animals (to reduce the level of potential secreted ascarosides) and found that their development was not restored (Figure S5S). Taken together, these data indicate C5 ascarosides (such as ascr#9 and icas#9), ∆C7 (ascr#7), ∆C9 (ascr#3) and C9(ascr#10) ascarosides likely act synergistically as hormones, or less importantly as pheromones, to negatively regulate animal development in GlcCer-deficient animals.

### Peroxisomal-derived ascarosides interact with sensory neurons to regulate development

We then investigated the downstream mechanism underlying the ascarosides dependent developmental repression. Most nematode ascarosides bind and activate their receptors located on the sensory cilia of chemosensory neurons(Inglis et al., 2007). If C5 and ∆C7 ascarosides regulate development by binding with their own sensory cilia, then disruption of the sensory cilia formation would also suppress the GlcCer deficiency-caused developmental arrest. CHE-3 is a homolog of human dynein cytoplasmic 2 heavy chain 1 and is essential for intraflagellar transport (IFT), a critical step for cilium biogenesis (Inglis et al., 2007). We found that *che-3(−)* also significantly rescued the development in GlcCer deficient animals (Figure 5D). In addition, in the normally developed *che-3(−); cgt-1/2/3(−)* animals, the peroxisomal apical accumulation was not reversed (Figure 5E, F). These data not only support our hypothesis that sensory neurons act downstream of peroxisomal ascaroside secretion, but also indicate a physiological role of intestinal apical accumulation of peroxisomes as a cause of the animal arrest under the GlcCer-deficient condition. Last, we found knocking down the nuclear hormone receptor DAF-12, a nuclear hormone receptor known to act downstream of ascaroside sensing that is essential for ascarosides-dependent dauer entry(Hu, 2007), could significantly suppress the GlcCer deficiency-induced developmental arrest (Figure 5G). The partial suppression suggests that additional regulatory proteins may also respond to the GlcCer/mTOR1/C5/∆C7 ascarosides dependent developmental regulation. Taken together, we conclude that peroxisomal beta-oxidation derived ascarosides are the negative developmental signal under GlcCer deficiency.

### Animals defective in GlcCer biosynthesis and PRX-11 function survived poorly under prolonged fasting

To learn more about the physiological functions of the GlcCer/peroxisome-mediated nutrient sensing system, we evaluated the lifespan of the *prx-11(−); cgt-1/2/3(−)* animals under well-fed and fasting conditions. While these animals displayed moderately decreased lifespan under the well-fed condition (Figure 5H), they survived poorly under prolonged fasting (Figure S5R). These data are consistent with the idea that the GlcCer lipid biosynthesis pathway and the peroxisomal derived hormones/pheromones are positive and negative regulatory machineries, respectively, that act coordinately with mTORC1 to enable the animal to adapt to fluctuated nutrient environment (Figure 6A,6B).

**Figure 6.**
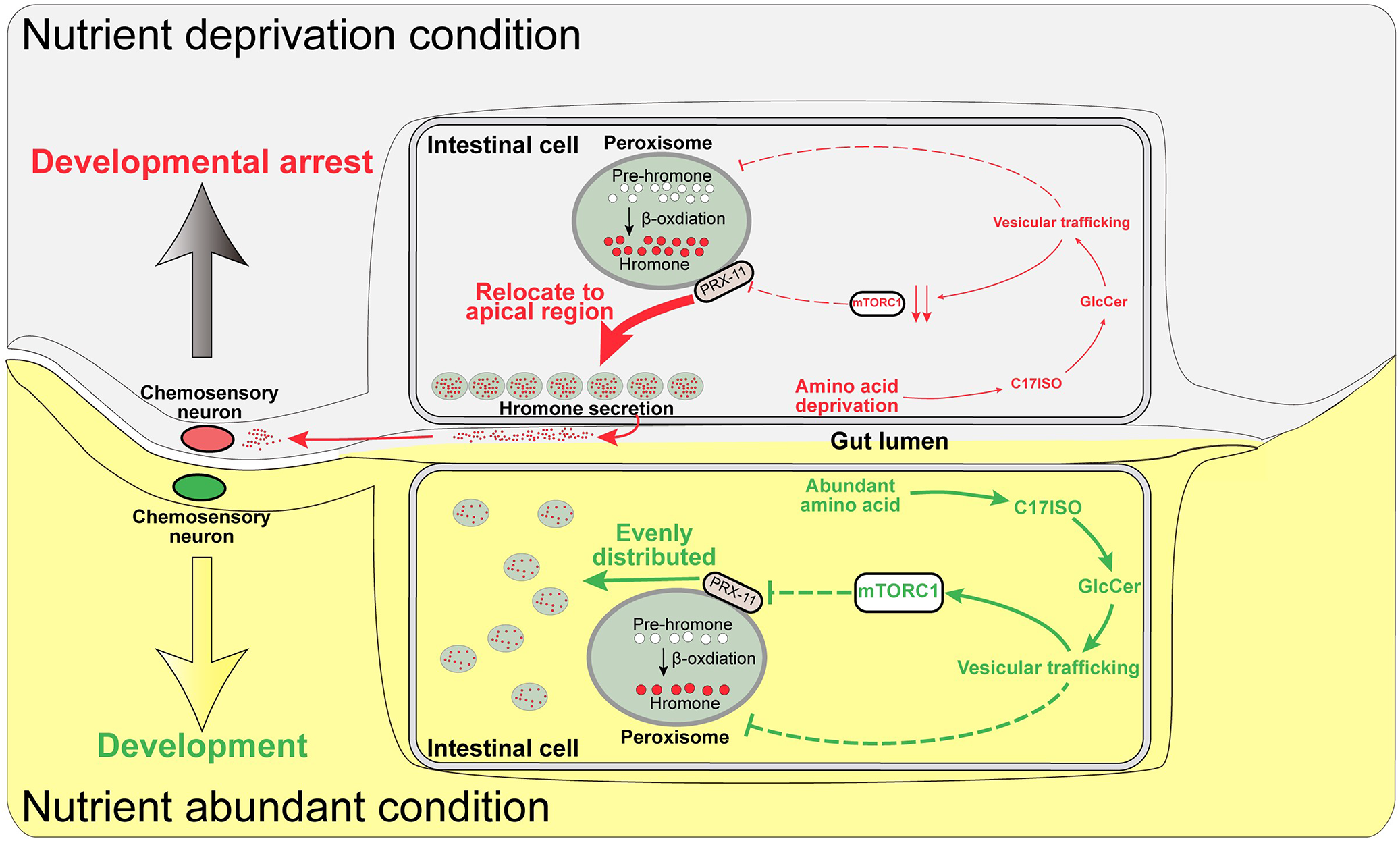
A model showing that GlcCer/mTORC1 negatively regulates the peroxisomal-derived ascarosides to promote *C. elegans* development.

## Discussion

### A new gut-brain axis regarding the roles of glycosphingolipid and peroxisomes in mTORC1-mediated nutrient sensing mechanism in animals

Among the diverse signaling functions of mTORC1, the mTORC1-mediated nutrient sensing machinery plays critical roles in orchestrating the early developmental process among metazoans(Guertin et al., 2006; Jia et al., 2004; Long et al., 2002; Oldham et al., 2000; Qi et al., 2017). Although extensive studies in the mTORC1 field in the past two decades have made major advances in revealing molecular mechanisms underlying the sensing of various nutrients, much of the work used cultured cells and the findings have not been clearly demonstrated *in vivo* under physiological conditions, including the mTORC1 role in regulating development. Besides the obscurity of the mechanisms underlying the connection between specific dietary nutrients such as AA and mTORC1 during development, the key downstream pathway that executes the mTORC1-mediated nutrient signal in animal development has remained elusive. Because sensing of dietary nutrients typically initiates in the intestine and involves cross-tissue communications, including the intestine-CNS talk to coordinate development, model organism-based assay systems need to play an even bigger role in exploring these mechanisms. Studies using model animals might have been slowed by numerous technical difficulties, such as withdrawing specific essential nutrient from food for live animals, separating nutrient sensing function of mTORC1 from other cellular functions, or defining roles of mTORC1 functions in specific tissues. Aided by the ability to deplete and supplement specific lipids and make tissue-specific genetic manipulations in *C. elegans*, we previously found that the core glycosphingolipid GlcCer, is necessary and sufficient to activate mTORC1 in the postembryonic development of *C. elegans,* and such a regulatory pathway is also conserved in mammalian tissue-cultured cells(Zhu and Han, 2014; Zhu et al., 2015; Zhu et al., 2013). Moreover, we have provided evidence that this obscure lipid pathway plays a key role in mediating the sensing of overall amino acids by mTORC1 under poor nutrient conditions and such a role is also likely conserved in mammals (Zhu et al., 2020). The data in this paper made another major advance by presenting a new gut-brain axis model (Figure 6) to explain how this lipid-mTORC1 pathway mediates the regulation of animal development by the availability of dietary nutrients and derived metabolites. In this regulatory response, the peroxisome plays a pivotal role by translocating to the intestinal apical region to produce and secrete “development-antagonizing” hormones/pheromones in response to the low intestinal lipid-mTORC1 activity.

### Peroxisome-derived hormone critically mediates the GlcCer-mTORC1 nutrient-sensing pathway for developmental control

For multicellular organisms such as *C. elegans*, it has been a long-time, puzzling question regarding how mTORC1 dependent nutrient signals in the intestine pass across the whole body to orchestrate developmental regulation, as the known activities downstream of mTORC1 signaling are expected to only function intracellularly (Blackwell et al., 2019). In this study, we found that short chain ascarosides, generated by peroxisome beta-oxidation and possibly secreted by peroxisomes under GlcCer/mTORC1 deficiency, negatively regulate postembryonic development, which is somewhat consistent with their well-known role in dauer diapause (promoting dauer entry and thus suspend larval development). Our data uncovered a new peroxisome route to facilitate mTORC1 dependent inter-tissue communication in addition to a previously described function in neuronal pathways(Uno et al., 2015). Such a mechanism also enables communication between animals in a population (Figure 5C) (aka as a pheromone). Although the peroxisome was known to produce steroid hormones in mammals, the production of hormones was often promoted by peroxisome proliferator-activated receptors under favorable nutrient conditions (Diano et al., 2011; Issemann and Green, 1990). However, in our study, we found the expression of key peroxisome beta-oxidation enzymes such as DAF-22 (which was thought to be critically upregulated to boost ascaroside biosynthesis under other stress conditions(Joo et al., 2010; Joo et al., 2009)) were not upregulated under GlcCer/mTORC1 deficiency or fasting condition. Alternatively, because we found that *C. elegans* peroxisomes contain intralumenal vesicles, which may store hydrophilic hormones/pheromones such as ascarosides, it is also possible that nutrient conditions regulate the secretion, instead of biosynthesis, of peroxisome derived hormones/pheromones.

One possible mechanism to control peroxisomal hormone release could be intestinal apical-translocation of peroxisomes under prolonged fasting or GlcCer-mTORC1 mediated nutrient sensing deficiency. The intracellular transportation of peroxisome was revealed more than 25 years ago(Rapp et al., 1996). In the years since, mostly in studies using *Drosophila* and mammalian cultured cell lines, scientists have found peroxisomal transportation moved along microtubules in a kinesin/dynein dependent manner, (Ally et al., 2009; Kural et al., 2005). However, it remains unclear what physiological factors affect peroxisomal distribution and what are the physiological consequences of different peroxisomal distribution patterns, especially under *in vivo* conditions(Covill-Cooke et al., 2021). Interestingly, a recent paper showed that some peroxisomes were translocated to lipid droplets and facilitated lipolysis in adult *C. elegans* and mouse adipocytes, which also suggests that the peroxisome translocation is needed under poor nutrient condition(Kong et al., 2020). It would be necessary to investigate further how the GlcCer/mTORC1 pathway regulates the peroxisomal translocation and whether such a mechanism under unfavorable nutrient conditions during development is conserved in mammals.

### Roles of GlcCer and peroxisome in lipid-mTORC1 mediated nutrient sensing mechanism during development may be the most critical part of their essential function in metazoans

GlcCer molecules are the core of the whole family of glucosylsphingolipid that are known to have many important physiological functions. We found GlcCer is essential mainly because it acts to repress peroxisomal derived hormone/pheromone secretion, which negatively regulate postembryonic development. Therefore, other reported roles of GlcCer, including being constitutes of many membrane structure such as lipid raft and being building blocks of the whole family of high-order glucosylated-glycosphingolipids that are associated with many important biomolecules(Merrill, 2011; Varki, 2017; Yoshihara et al., 2018), are not essential and might be recruited to the eukaryotes relative later during the evolution. This notion is consistent with the idea that, although the origin of glycosphingolipid and related enzymes are relative early (some bacteria already produce GlcCer(Kawahara et al., 2001)), it is not essential in bacteria and many unicellular eukaryotes such as budding yeast(Sibirny, 2016), but absolutely essential in multicellular metazoans from *C. elegans* to human(Kohyama-Koganeya et al., 2004; Marza et al., 2009; Monies et al., 2018; Yamashita et al., 1999). We hypothesize that GlcCer was recruited to mediate the mTOR dependent nutrient sensing in multicellular metazoans because, unlike in unicellular eukaryotes, it is much more critical for the multicellular metazoans to efficiently monitor nutrient levels, especially those that cannot be biosynthesized endogenously (such as essential amino acids). This concept was confirmed in our recent finding that mmBCFA-derived GlcCer acts as a key signal for overall AA abundance to promote *C. elegans* larval development even under poor AA condition(Zhu et al., 2020). Given that both blocking GlcCer biosynthesis and mutating the major mTORC1 components caused murine embryos to die quickly after implantation(Guertin et al., 2006), it would be interesting to investigate whether the similarity in phenotypes is also due to disruption of a conserved GlcCer/mTORC1 nutrient sensing machinery.

Similarly, it is also interesting to speculate why a “housekeeping” organelle like the peroxisome is involved in the GlcCer/mTORC1-dependent developmental regulation. Like GlcCer, the peroxisome is ubiquitously present in eukaryotes but not essential for the growth of unicellular eukaryotes *S. cerevisiae*, at least under favorable nutrient conditions(Sibirny, 2016). Under other nutrient conditions (such as fatty acid or methanol as the exclusive carbon source), yeast cannot grow without functional peroxisomes, possibly due to “toxicity accumulation” (Sibirny, 2016). In addition, works in yeast indicated that peroxisome-depleted yeast could regenerate peroxisomes from mitochondria and ER derived vesicles(Yuan et al., 2016), which suggests its late origin in the evolution. However, in metazoans, the peroxisome is absolutely required for normal development and deficiency will cause severe developmental defects. For example, disruption of multiple *C. elegans* peroxins causes early developmental arrest(Thieringer et al., 2003). In humans, peroxisome disorders are a large family of developmental disorders due to genetic mutations in peroxisomal biogenesis (PBD) and peroxisomal enzyme. PBD patients suffer from severe developmental defects in neuronal tissues and in other organs and may die in utero or in the first year after birth(Argyriou et al., 2016). Such prominent differences between unicellular and multicellular eukaryotes suggests that, like GlcCer, the essential roles of peroxisome in higher organisms could be gradually adapted during evolution, though the exact reason is not clear.

Taken together, the two “lately adopted machinery” GlcCer and peroxisome, had potentially integrated together and established a “check-and-balance” system in *C. elegans* to regulate the development. While GlcCer may be recruited as a critical metabolite to positively facilitate the mTORC1 dependent nutrient sensing pathway, the peroxisome may clear harmful metabolites(Govindan et al., 2019; Zhang et al., 2019) (deficiency of peroxisomal beta oxidation in humans and mice showed brain and liver pathology(Islinger et al., 2018)) to sustain normal development under nutrient abundance, or deliver a negative signal to slowdown or halt the development under unfavorable nutrient condition. The ability of the peroxisome to biosynthesize small metabolic hormone/pheromone could be ideal for such a negative regulatory role. Therefore, a multicellular organism equipped with these two machineries (GlcCer and peroxisome) may better adapt to the fluctuating nutritional environment in nature, as we showed in this work (Figure 6).

## Acknowledgement

We thank Shohei Mitani, Chenggang Zou, Shiqing Cai, Di Chen, Hongyun Tang, Xiajing Tong, Guisheng Zhong, Shengjie Fan, Cheng Huang, Szecheng J. Lo and the *C. elegans* Knockout Consortium and CGC (funded by NIH [P40OD010440]) for strains, reagents and advice. We thank all of our laboratory members for helpful discussions. We thank Ying Han, Piliang Hao from the Multi-Omics Core Facility (MOCF), Xiaoming Li, Ziwei Yang and Chengyu Fan from the Molecular Imaging Core Facility (MICF), Ying Xiong, Xiaoyue Ren from the Molecular and Cell Biology Core Facility (MCBCF) at the School of Life Science and Technology, ShanghaiTech University, Lei Zhang from Professional Technical Support Sharing Platform of Core facility (Improvement of Service Capability for Shanghai Proteomics Professional Technical Platform for Severe Diseases [grant number 18DZ2292900]) of Shanghai Medical College at Fudan University for providing technical support. Funding: This work was supported by the Recruitment Program of Global Experts of China (Youth), the Shanghai Pujiang Program (16PJ1407400), the National Key R&D Program of China (2019YFA0802804) and ShanghaiTech Startup program. Author contributions: NL, Conception and design, Acquisition of data, Analysis and interpretation of data, NL, HZ Drafting or revising the article; NL, QC, MR, MZ, HS, LZ Acquisition of data, Analysis and interpretation of data, Drafting or revising the article; Contributed unpublished essential data or reagents; HZ Supervised the study, Conception and design, Drafting or revising the article. Competing interests: Authors declare no competing interests. Data and materials availability: All data is available in the main text or the extended data materials.

