## supplementary figures and legends for "A lipid-mTORC1 nutrient sensing pathway regulates animal development by peroxisome-derived hormones"

**Figure S1**

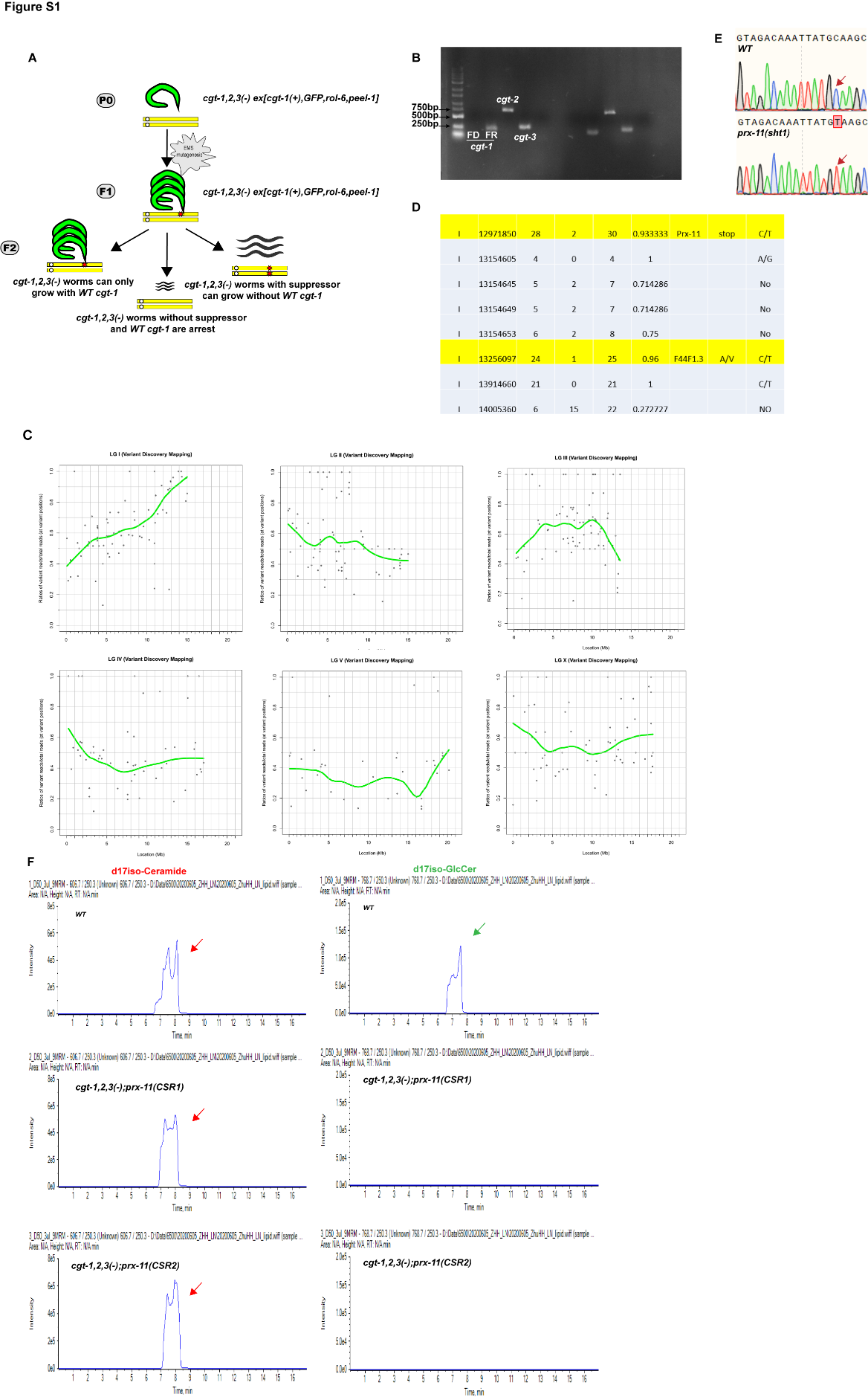

**Figure S1. *prx-11* mutants render GlcCer dispensable in *C. elegans* development, Related to Figure 1.**

(A)**.** The scheme of genetic suppressor screen for mutants that suppressing GlcCer deficiency-induced L1 arrest. We subcloned the WT *cgt-1* (with its promoter and the coding region) and made a *WT-cgt-1* transgenic line in *cgt-1/2/3(-)* with and three selection markers *sur-5::GFP*, *hsp-16_prom_peel-1* and *rol-6* genes. This *cgt-1/2/3; ex[WT-cgt-1]* strain was viable, fertile, and seemed healthy (Figure. 1B), confirmed the previous finding that *cgt-1* overexpression could largely compensate the deficiency of endogenous GlcCer synthase(Marza, Simonsen et al. 2009), and enable us to perform the following genetic suppressor screen. Normally the *WT-cgt-1* carrying extrachromosomal array may be automatically lost in the offspring by chance, and such animals would arrest at the early larval stage. However, if a recessive mutant could suppress the *cgt-1/2/3* caused the developmental arrest and grew to adulthood in the F2 generation, they could be easily identified by loss of both positive (*sur-5::GFP* and *rol-6*) and the negative (*hsp-16_prom_peel-1*) selection markers (Figure. 1F). (B) PCR experiments confirmed that no *WT-cgt-1* sequence was in the *cgt-1/2/3; prx-11 (sht1)* animals. (C) Dot/Curve graphs showing the penetrance of EMS-generated mutations in *prx-11(sht1)* animals. Briefly, we outcrossed *cgt-1/2/3; prx-11(sht1) to cgt-1/2/3; ex[WT cgt-1]* strain (the original strain before EMS mutagenesis) and pooled the F2 adults for deep sequencing. *prx-11(sht1)* clearly located at the far-right part of chromosome I. (D) A table showing all mutations at the mapped locus. Mutations that changed the protein sequence were marked in yellow. (E) DNA sequence graphs showing that a C-T transition in the gene *prx-11*. (F) Multiple Reaction Monitoring (MRM) graphs by UPLC-Mass spectrometry showing peaks of ceramide (red/left) and GlcCer (green/right) in *cgt-1/2/3; prx-11(-)* mutants.

**Figure S4**

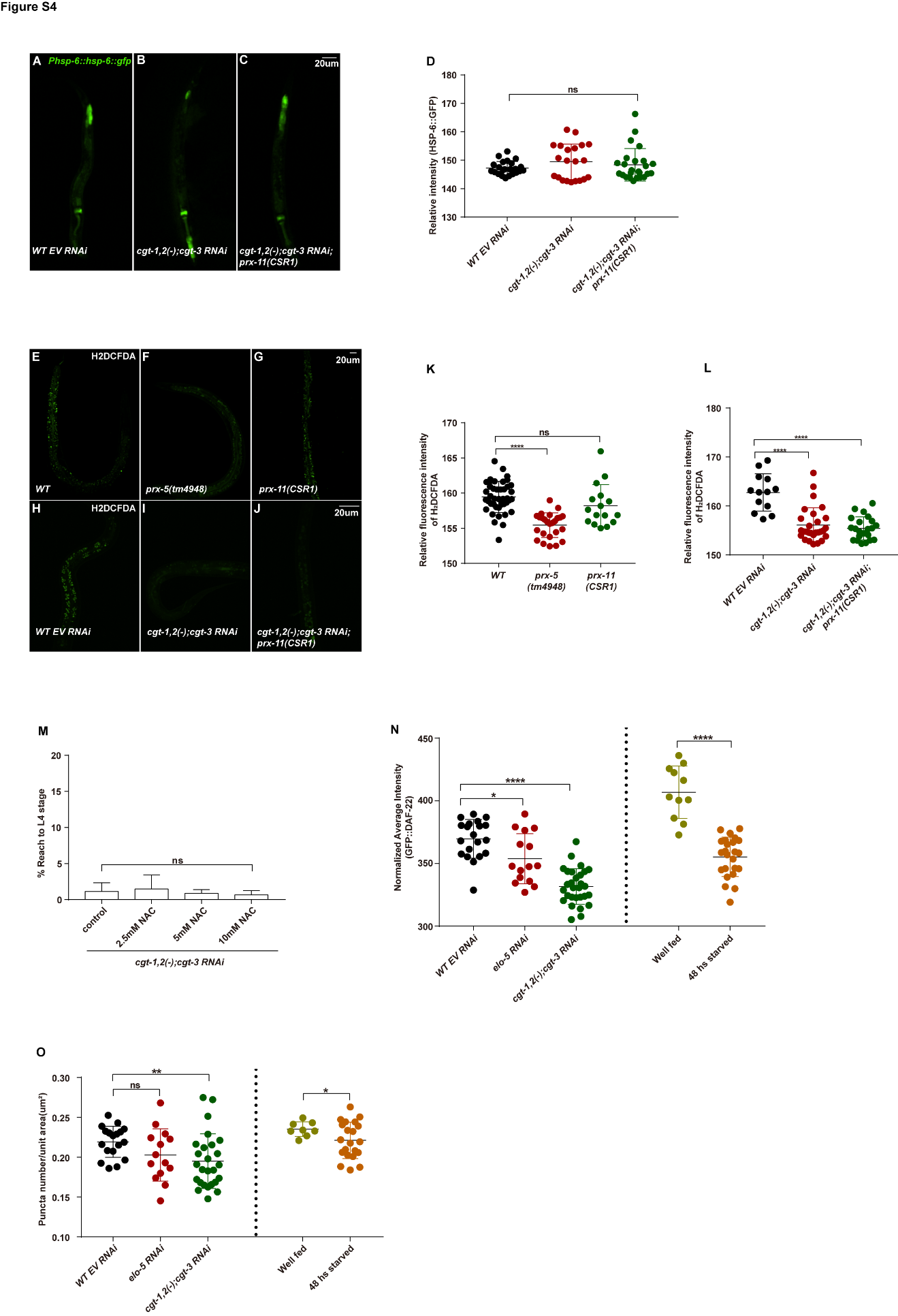

**Figure S4. Peroxisome deficiency did not suppress GlcCer deficiency induced developmental arrest by increased ROS or general peroxisomal activity, Related to Figure 4.**

(A-D) Representative fluorescent microscopic images (A-C) and statistics (D) showing that the intensity of HSP-6::GFP in L1 animals with indicated genotypes. Neither glucosylceramide deficiency (B) nor *prx-11(-)* (C) affected the HSP-6::GFP expression. (E-L). Representative fluorescent microscopic images (E-J) and statistics (K, L) showing that the intensity of H2DCFDA (a marker of ROS) in L1 animals with indicated genotypes. The ROS level was reduced in *prx-5(tm4948)* (F) and glucosylceramide deficient animals (I), and *prx-11(-)* could not further decrease the ROS activity in glucosylceramide deficient animals (J). (M) A bar graph showing the percentage of GlcCer deficient animals grew beyond L4 stage under the treatment of ROS inhibitor N-acetyl-l-cysteine(NAC). NAC could not suppress glucosylceramide-induced developmental arrest at various concentrations. (N, O). Dotted graphs showing numbers of peroxisomes(O) and their activity(N) (measured by the level of DAF-22) under indicated genotypes and food conditions. Neither numbers of peroxisomes nor peroxisomal activity were increased under *elo-5* RNAi, GlcCer depletion, or starvation. Data are represented as mean ± SD.

**Figure S5**

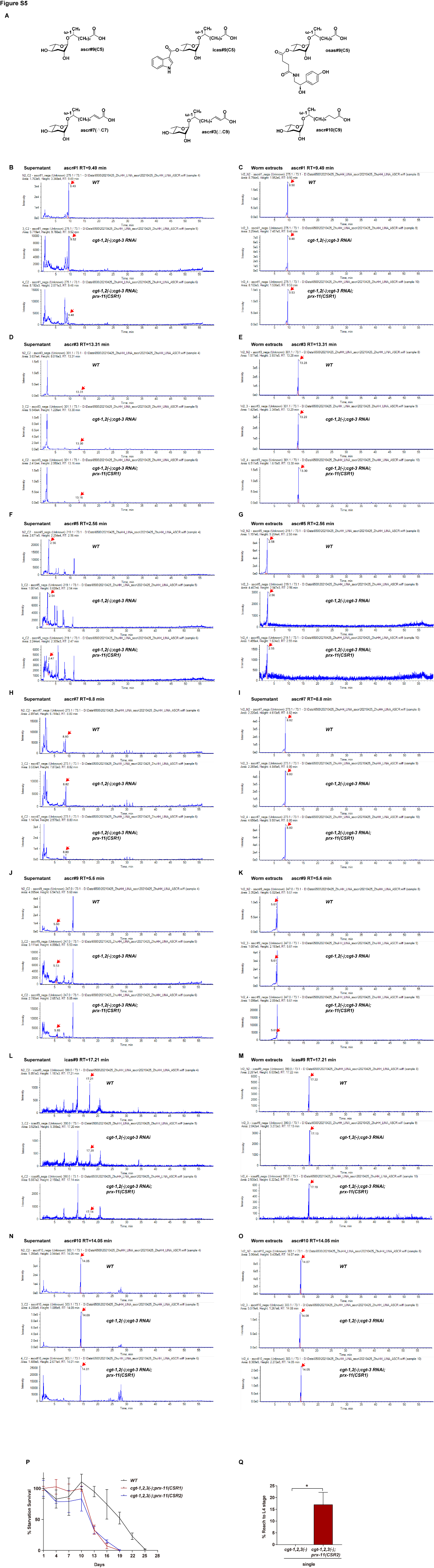

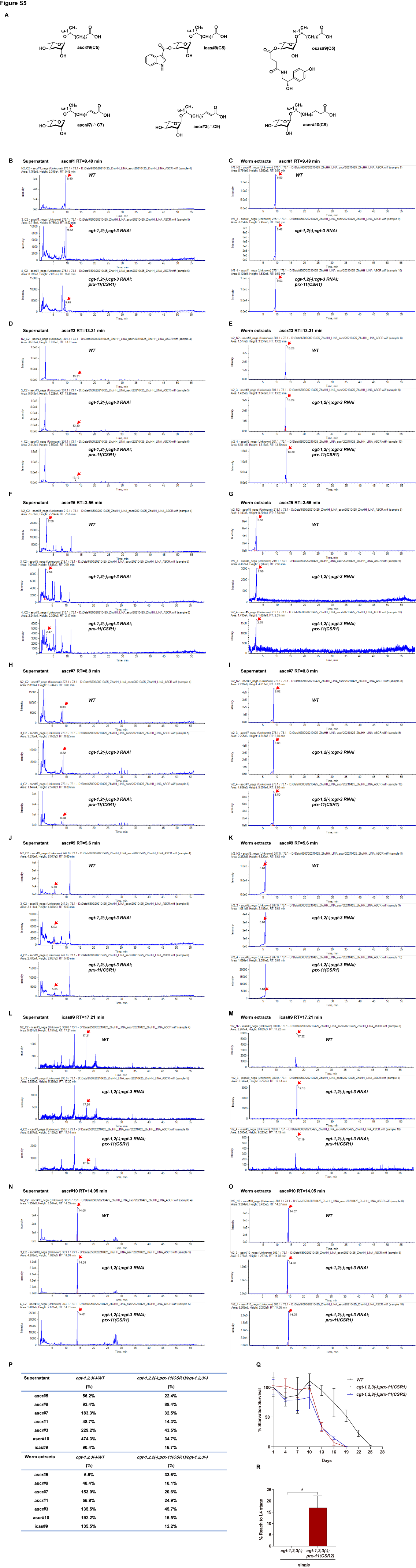

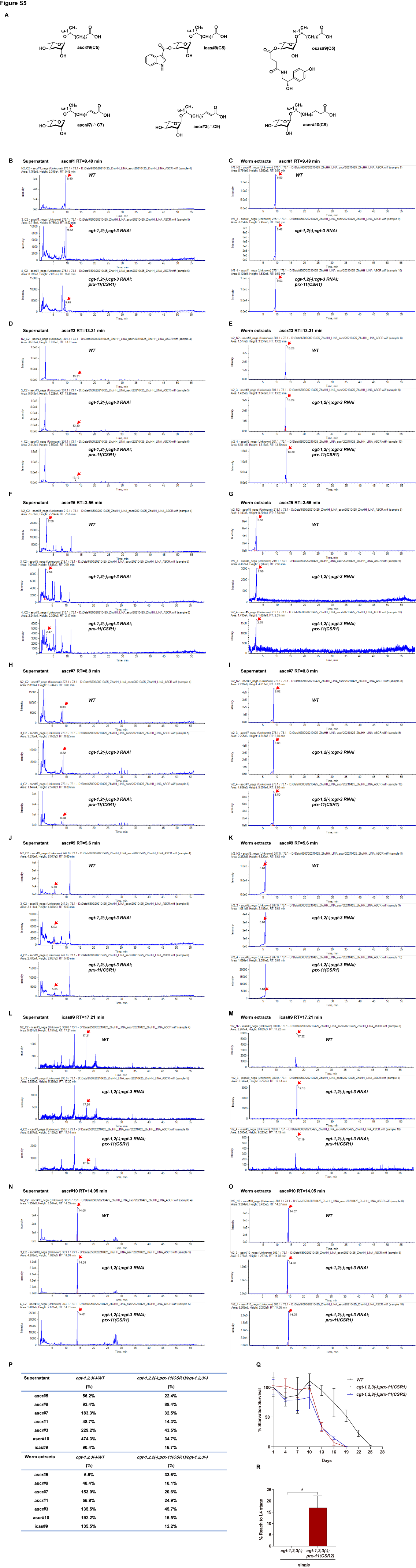

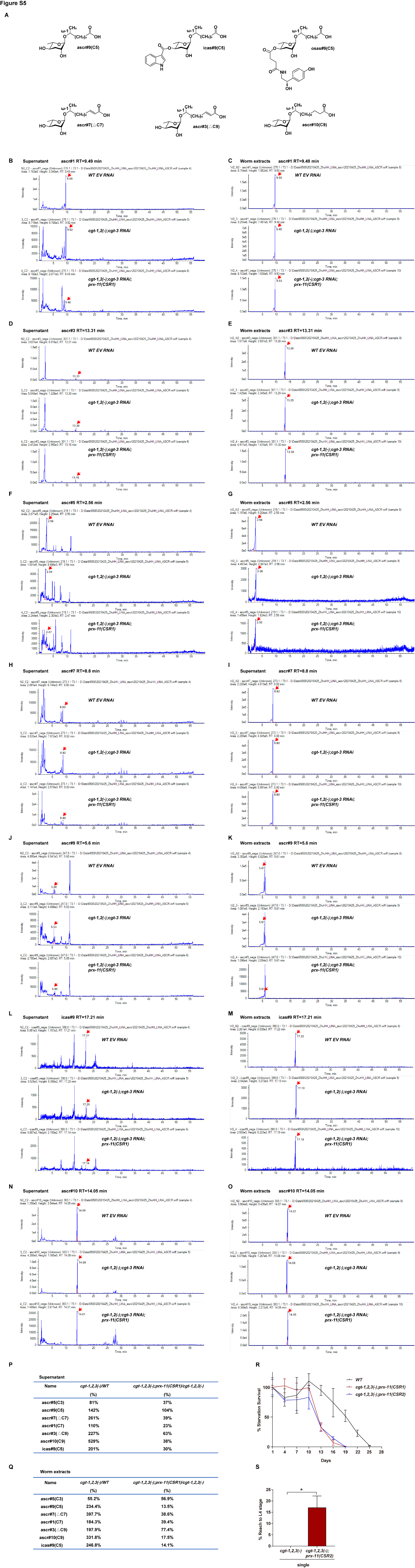

**Figure S5**. **Peroxisomal beta-oxidation derived ascarosides repress GlcCer/mTOR promoted development, Related to Figure 5.**

(A) Chemical structures of C5, ΔC7, ΔC9 and C9 ascaroside. (B-L) Mass spectrometry by precursor scan m/z 73.1 showing ascaroside standards of *WT EV RNAi, cgt-1,2(-);cgt-3 RNAi and cgt-1,2(-);cgt-3 RNAi;prx-11(CSR1)* in supernatant of culture media(B,D,F,H,J,L,M) and worm extracts (C,E,G,I,K,M,O). (P, Q) Table of relative peak intensity ratio from supernatant extracts (P) and worm body extracts (Q) by UPLC-Mass spectrometry showing several ascarosides (ascr#7, ascr#9, icas#9 and ascr#10) were increased in *cgt-1,2(-);cgt-3 RNAi* compared to WT and decreased in *cgt-1/2/3;prx-11(-)* mutants compared to *cgt-1,2(-);cgt-3 RNAi* . (R) Bar graphs showing percentage of animal grew beyond L4 stage under various growth conditions. (S) Survival curves showing that GlcCer depleted animals survived poorly under prolonged starvation. Data are represented as mean ± SD.
