## Supplementary material for "A lipid-mTORC1 nutrient sensing pathway regulates animal development by peroxisome-derived hormones": Materials and methods

**STAR★METHOD**

**KEY RESOURCES TABLE**

| REAGENT or RESOUCES | | SOURCE | IDENTIFIER |
| --- | --- | --- | --- |
| Antibodies | | | |
| Fibrillarin antibody[38F3] –  Nucleolar Marker | | abcam | Cat#ab4566; RRID:AB_304523 |
| Goat anti-Mouse IgG(H+L)  Cross-Adsorbed Secondary  Antibody, Alexa Fluor 594 | | Thermo Fisher Scientific | Cat#A-11005; RRID:AB_2534073 |
| Chemicals, Peptides, and Recombinant Proteins | | | |
| DAPI Staining Solution | | beyotime | Cat#C1005 |
| N-Acetyl-L-cysteine | | Sinopharm | Cat#62024261 |
| N-butyroyl-D-erythro-sphingosine | | aladdin | Cat#74713-58-9 |
| Bacteria | | | |
| OP50 | | CGC | https://cgc.umn.edu/strain/HT115(DE3) |
| OP50 RNAi | | Jianfeng Liu’s Lab | N/A |
| BL21 | | Jianfeng Liu’s Lab | N/A |
| Experimental Models: Oganisms/Strains | | | |
| *C. elegans* Strains  see Table S1 | | This paper | N/A |
| Recombinant DNA | | | |
| Plasmid: *Pprx-11::prx-11::gfp* | | This paper | N/A |
| Plasmid: *Prgef-1:prx-11* | | This paper | N/A |
| Plasmid: *Pges-1::prx-11* | | This paper | N/A |
| Plasmid: *Pdpy-7::prx-11* | | This paper | N/A |
| Plasmid: *Pmyo-3::prx-11* | | This paper | N/A |
| Plasmid:*Ppmp-2::pmp-2::mCherry* | | This paper | N/A |
| Plasmid: *Phsp-16.2::gfp-SKL* | This paper | | N/A |
| Software and Algorithms | | | |
| GraphPad Prism 7 | | GraphPad Software | https://www.graphpad.com/scientificsoftware/prism/ |
| ImageJ | | NIH | https://imagej.nih.gov/ij/ |

**RESOURCE AVALABILITY**

**Lead Contact**

**Materials Availability**

All *C. elegans* strains and plasimds generated in this study are available on request from the Lead Contact.

**Data and Code Availability**

There is no database or code generated from this paper.

**EXPERIMANTAL MODEL AND SUBJECT DETAILS**

**Animal Models**

*C. elegans* were maintained at 20°C on NGM plates (referred to as standard plates) with *E. coli* OP50 as the bacterial food (OP50/NGM). Details and a complete list of strains in this study are shown in Table S1.

**Materials and methods**

***Caenorhabditis elegans* strains and maintenance**

All worm strains were cultured under standard conditions(Brenner 1974). The bleaching process was according to the standard protocol(Stiernagle 2006). The following strains were obtained from the *Caenorhabditis* Genetics Center Database (CGC) or as indicated; wild type N2 Bristol, *raga-1(ok386)*, *prx-5(ku517)*, *bre-3(ye26)*, *prx-11(gk959960)*, *che-3(e1124)*, *daf-12(rh61rh412)*, *osm-3(p802)*. The *cgt-1(tm1027)*, *cgt-2(tm1192)*, *cgt-3(tm504)* and *prx-5(tm4948)* mutants were provided by the Mitani Lab (National BioResource Project, Tokyo, Japan).

**Isolation of *prx-11(sht1)* through a genetic screen -EMS mutagenesis**

We subcloned the WT *cgt-1* (with its promoter and the coding region) into the *C. elegans* expression vector pPD95.77, and microinjected the plasmid into the *cgt-1/2/3(-)* animals with three selection markers *sur-5::GFP, hsp-16_prom_peel-1*, and *rol-6* genes. This *cgt-1/2/3(-); ex[WT-cgt-1]* strain was viable, fertile, and seemed healthy.

A Genetic screen of GlcCer depletion suppressors

The EMS mutagenesis on the *cgt-1/2/3(-); ex[WT-cgt-1]* strain was performed as previously described(Zhu, Shen et al. 2013). We have obtained 49 mutants that could survive without carrying the positive selection markers (roller and GFP)*.* PCR experiments were performed to verify the *WT-cgt-1* was no longer exist.

Genetic mapping of *prx-11(sht1)*

We used genetic linkage analysis to map the *prx-11(sht)* mutation (zhu 2013 eLife)*,* we outcross *cgt-1/2/3(-); prx (sht1)* to *cgt-1/2/3(-);ex[WT cgt-1]* strain (the original strain before EMS mutagenesis) and pooled the F2 adults. Then we extracted the genomic DNA of those adults using the Genomic DNA Sample Preparation Kit (Illumina) and send for Illumina deep sequencing (High Throughput Next-Generation Sequencing Core, Novogene company). We compared the genome sequences of *prx-11(sht1)* animals and the *ex[WT-cgt-1]* animals by using CloudMap Workflows(Doitsidou, Jarriault et al. 2016). All candidate mutations were confirmed by PCR and Sanger sequencing.

**Transgenic animals**

For *Ppmp-2::pmp-2::mCherry*, the genomic DNA including the full coding region and about 3 kbps of the upstream sequence was cloned into pPD95.75 vector. For *Pprx-11::prx-11::gfp*, the genomic DNA including the full coding region and about 3 kbps of the upstream sequence was cloned into pPD95.77 vector. For *ges-1::prx-11*, *rgef-1::prx-11*, *myo-3::prx-11,* and *dpy-7::prx-11*, the genomic *prx-11* DNA including the full coding region was cloned into pPD95.77, driven by the indicated promotors respectively. All plasmids were extracted by QIAGEN kit and injected to indicated worms.

**Genome Integration of Extrachromosomal Arrays Transgenic animals**

TMP/UV Integration was performed as previously described (Kage-Nakadai, Imae et al. 2014). The *Ppmp-2::pmp2::mCherry* and *Phsp-16.2::gfp-SKL* plasmids were integrated into the genome of *C. elegans* through TMP/UV treatment.

**Generation of Knock-in/out animals**

The CRISPR Cas-9 associated knock-out/in experiments was according to the previous articles (Dickinson, Ward et al. 2013, Shen, Zhang et al. 2014, Paix, Folkmann et al. 2015). For the Knock-out experiments, we designed sgRNA of *prx-11 (CSR1 and CSR2)*(sgRNA1: 5’-cagtactctcttcatatgc-3’; sgRNA2: 5’-atcttagcgtcactaagcc-3’) and constructed Cas9‐sgRNA plasmids. The plasmids were purified, mixed, and injected to indicated genotype worms. For the Knock-in experiments, we designed sgRNA of *prx-11(sht1)*(sgRNA 5’-cttttcaaactggaagaat-3’) and *daf-22*(sgRNA1: 5’-atgacgccaaccaagccaa-3’; sgRNA2: tgtatacctttggcttggt) and constructed Cas9‐sgRNA plasmids. For *gfp-daf-22* knock-in, a gfp coding sequence flanked by 40bp homologous arms of *daf-22* coding region and the *daf-22* promoter for homologous recombination was constructed. Then the sgRNA plasmids and the repaired template were mixed and injected into worms with indicated genotypes.

**Peroxisomal number and intensity test**

The measurement of peroxisomal numbers and intensity was performed as previously described (Narayan, Ly et al. 2016). For imaging of *gfp-daf-22* knock-in animals, L4 worms were anesthetized on 2% agarose pads and subjected to Nikon CSU SORA spinning disk microscope. The puncta numbers and intensity of peroxisome were analyzed using Fiji software. For *hsp-16.2*-GFP-SKL animals, they were shifted to 34℃ for 1.5h, then returned to 20℃ for 12h before being imaged.

**Assay for ROS activities**

Direct measurement of intracellular ROS assay

The measurement of ROS assay was performed as previously described (Coppa, Guha et al. 2020). Briefly, the 2,7-dichlorodihydrofluorescein-diacetate (H_2_DCFDA), a membrane-permeable non-fluorescent dye, was used to measure the ROS activity. L4/L1 stage worms were collected and washed three times with M9 buffer to remove the bacteria. Then worms were transferred into the staining solution (H_2_DCFDA 10uM in M9 buffer) and stained for 30 min at 20℃ before washed twice with M9 buffer and mounted to an agarose pad and subjected to Nikon CSU SORA spinning disk microscope.

HSP-6::GFP expression test.

Excessive reactive oxygen species (ROS) can lead to mitochondrial unfold protein response (mt UPR) and ready to be detected by HSP-6, a normally used marker of mt UPR activation in *C. elegans* (Runkel, Liu et al. 2013). For the *WT EV RNAi*, *cgt-1,2(-);cgt-3 RNAi* and *cgt-1,2(-);cgt-3 RNAi;prx-11(CSR1)* animals, *C. elegans* were grown at the indicated developmental stages and HSP6::GFP fluorescence signals were determined by Nikon CSU SORA spinning disk microscope.

ROS inhibitor (NAC) treatment assay

The NAC treatment assay was modified and performed as previously (Yang and Hekimi 2010). Briefly, the NAC stock solution (50mM in DMSO) was added into *cgt-3 RNAi* bacteria medium into the indicated concentrations and seeded on NGM plates. Then L4 stage *cgt-1,2(-)* worms were transferred to those *cgt-3 RNAi* plates and their progenies beyond L4 stage were scored.

**mTORC1 activity assay**

FIB-1 staining assay

The immunofluorescence assay of FIB-1 was done according to a previously published “frozen crack” method (Duerr 2013). Briefly, L1 stage worms were collected and washed three times with distilled water to remove bacteria. Then the worms were adhered to slides coated with polylysine and “frozen crack” was used to expose the intestine. Subsequently, the worms were fixed with 4% Paraformaldehyde (PFA) for 30 min at room temperature(RT) immediately followed by three washes with PBS at RT. The samples were then blocked with 0.5% Normal Donkey Serum in PBS with 0.1% Triton-100(PBST) for 1 hour at RT. Primary and secondary antibody incubation was done in 0.5% BSA in PBST using Fibrillarin (Abcam ab4566,1:400) as the primary antibody and goat anti-mouse conjugated to Alexa Fluor 594(Invitrogen A11005, 1:800) as the secondary antibody. DAPI was applied for staining nuclei. Samples were washed with PBST after antibody treatments and finally mounted with Antifade Mounting Medium (Beyotime P0126). The imaging and quantification of FIB-1 were based on the previously article (Zhu, Shen et al. 2013, Tiku, Jain et al. 2017). Briefly, L1 stage worms were imaged using a Nikon CSU SORA spinning disk microscope. Then the intestinal cell nuclear (DAPI) and nucleolar (FIB-1) were quantified using Fiji software. Subsequently, the ratio of nucleolar/nuclear area was calculated.

The quantification of HLH-30::GFP

The imaging and quantification of HLH-30 were based on the previously article (Lapierre, De Magalhaes Filho et al. 2013). Briefly, Worms carrying HLH-30::GFP transgene were imaged using a Nikon CSU SORA spinning disk microscope. Then the nuclear localization of HLH-30::GFP in the intestinal cell was Quantified using Fiji software.

**RNAi feeding assay**

Because GlcCer deficient animals grew poorly on *E. coli* HT115 bacteria, all RNAi carrier bacteria strain was *E. coli* OP50 instead (Xiao, Chun et al. 2015) in this study, except specifically mentioned. Briefly, we extracted the dsRNA-expression plasmids from the HT115 host strain and then transformed it into the OP50 strain. The RNAi-feeding experiments were done as previously described(Kamath, Martinez-Campos et al. 2001). dsRNA-expression constructs were from our previous works (Zhu, Sewell et al. 2015)*(daf-15,let-363,cgt-3)*, from the ORF-RNAi library (Open Biosystems)(*aps-1,dhs-28,daf-22,acs-13,acox-3,acs-7,daf-12*) or from the Ahringer RNAi library(Kamath, Fraser et al. 2003)(*prx-1, prx-3, fath-1, elo-5, sptl-1, chc-1, maoc-1, acox-1.1, acox-1.4*). *daf-15* RNAi bacteria were diluted with empty vector-containing bacteria to reduce the RNAi knockdown effect. In all RNAi experiments except *elo-5* RNAi, P0 worms of L4 stage (P0 L1 stage worms were used in *elo-5* RNAi experiment) were seeded on the RNAi plates.

**Bt toxin (Cry-5B) assay**

The toxicity assay was done as previously described(Marroquin, Elyassnia et al. 2000). Briefly, Bt-toxin (Cry-5B) expressing *E. coli* BL21 was cultured overnight in LB containing 50ug/ml kanamycin. Then bacteria was concentrated twice by centrifuge and seeded on plates(containing 50 ug/mL kanamycin, 1mM IPTG), beforeL4 stage worms with indicated genotype were added. Worm’s survival rate were recorded on day 4.

**Lipid analysis by UPLC/QTRAP-MS**

For sphingolipid analyses

Total lipids were extracted from about 0.2g worms, using a modified method of (Bligh and Dyer 1959)(Zhu eLife 2013). Lipid analysis is based on the previous article(Lin, Ding et al. 2017). Briefly, UPLC-MS/MS profiling was performed using SHIMADZU LC30A UHPLC system equipped with a Phenomenex Luna3μm silica column (100Å LC column 150*2mm) coupled to the SCIEX Qtrap6500. A gradient from solvent A (isopropanol/hexane/100mM ammonium acetate 58/40/2; v/v) to solvent B (isopropanol/hexane/100mM ammonium acetate 50/40/10; v/v) was used, starting with 50% B for 5 min which was increased to 100% from 5-30 min and remained for 10 min. B was decreased to 50% from 40-41 min which was remained until the end of the run at 50 min. The precursor scan m/z=+250.3 (CE=46) was performed using a positive ionization mode. Three independent experimental repeats were performed and the relative amount of Ceramide and GlcCer in WT and *cgt-1,2,3(-);prx-11(-)* was normalized by the level of internal standard d18:1/C4-Ceramide.

For ascaroside analyses, the metabolite extracts were analyzed based on the previously article(von Reuss, Bose et al. 2012, Qian, Zeng et al. 2021). Briefly, wild-type(*N2 L4440 RNAi*), triple mutant(*cgt-1,2(-);cgt-3 RNAi*) and quadruple mutant(*cgt-1,2(-);cgt-3 RNAi;prx-11(CSR1)*) worms were cultured on 90 mm plates(50 plates for each strain). Then the plates were washed at L1 stage using M9 buffer and triple/quadruple mutant was collected 24H later than WT to improve the RNAi efficiency. The supernatant media and worm pellets were evaporated in a vacuum at room temperature. Recorded the weight for each strain to correct the mass spectrum result. Then the supernatant media were extracted with methanol and 100ul extracts were injected for UPLC-MS analysis. The worm pellets were crushed with 0.2g NaCl using a mortar pestle and extracted with 100% ethanol at room temperature overnight. Then the suspensions were centrifuged and evaporated in a vacuum. The worm pellet metabolite extracts were remelted with methanol and 100ul extracts were injected for UPLC-MS analysis. UPLC-MS was performed using a Sciex TripleTOF 6500 system equipped with a Waters Acquity UPLC BEH C18 column (1.7 mm, 2.1*10 mm), with a flow rate of 0.4 ml/min. A gradient from solvent A (0.05% formic acid in H_2_O) to solvent B (0.05% formic acid in acetonitrile) was used, starting with 3% B for 1 min which was increased to 100% from 1-55 min. B was decreased to 3% from 55-55.1 min which was remained until the end of the run at 60 min. The precursor scan m/z=+73.1 was performed using a negative ionization mode. Data acquisition and processing were performed by Analyst and Peakview (Sciex).

**Analyses of the peroxisome intracellular distribution**

For *WT EV RNAi, cgt-1,2(-);cgt-3 RNAi, cgt-1,2(-);cgt-3 RNAi;prx-11(CSR1), aps-1 RNAi, chc-1 RNAi, daf-15 RNAi* and *let-3 RNAi* animals, *C. elegans* were grown at the indicated developmental stages. For starved animals, *C. elegans* were bleached and synchronized L1 were suspended in the M9 medium for 48 hours. The intracellular distribution of peroxisomes in the intestinal cells was documented by a Nikon CSU SORA spinning disk microscope. Pictures were then analyzed by Icy software. For each worm, a rectangle area which covered the half of intestinal cells was selected and the peroxisome- distribution-value was measured by the average value of individual peroxisome’s relative position X between the apical and the basal membrane (Peroxisomal distribution value was considered 0 at the apical membrane region, and 1 at the basal membrane region)(See pictures below). Therefore, a smaller value indicated that peroxisomes were more close to the apical region.

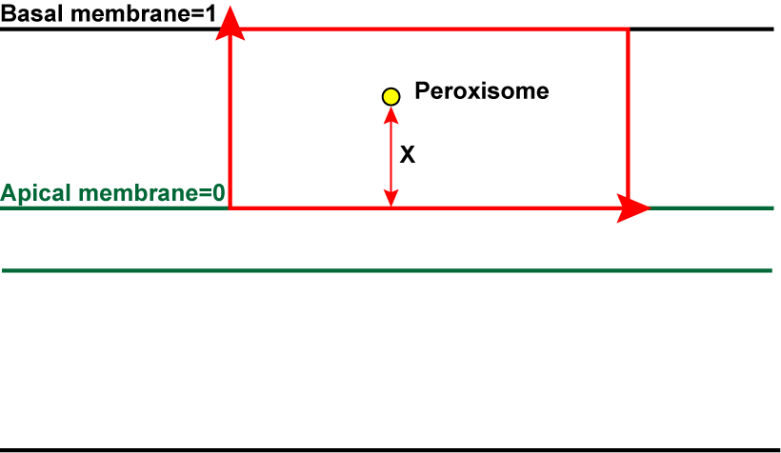

**Plate Assay for developmental-suppressing hormones**

The pre-incubation assay was similar to a previous article(Ludewig, Gimond et al. 2017). First, 4 animals (P0) of the indicated genotype were seeded on the *cgt-3* RNAi plates to lay eggs for two days before removed. These plates were considered pre-conditioned since a large number of F1 animals grew on the plates and potentially secreted ascarosides (For plates pre-conditioned by WT animals, we removed all F1 animals before they reached L4 stage to prevent the depletion of bacteria food). Then fluorescent-labeled *cgt-1/2/3(-); prx-11(-)* L1 animals were transferred into these pre-conditioned plates, 5 worms per plate. The number of worms that grew beyond L4 stage were counted after day 6.

**Lifespan assay**

The lifespan assay was done as previously reported method (Yin, Liu et al. 2014). Briefly, synchronized L1 worms were seeded on standard NGM plates at 20°C. Then L4 stage worms were transferred to plates as day 0(~30 worms per plate). Worms not responding to the repeated prodding were considered as dead. The number of the dead worms was recorded in every other day until all worms had died.

**L1 starvation survival assay**

The L1 starvation assay was done following the previous protocol (Cui, Cohen et al. 2013). We compared the animals’ survival rates in different genotypes by simulating the survival rate of each genotype to 100 arbitrary “individual worms”. The significance of the difference in overall survival rate was calculated by the log-rank test.

**Microscopy**

All *C. elegans* fluorescence microscopy images(except Fig 4 L-W) were acquired using a Nikon CSU SORA spinning disk microscope and a prime 95B photometric sCMOS camera. STED images of *C. elegans*(Fig 4 L-W) were captured using a Leica SP8 STED Microscope, with hybrid photon-counting detectors (HyD). Plate phenotypes were observed with an MVX-ZB10 fluorescence dissecting microscope and a BioHD-C20 CMOS camera. Fluorescence intensity and puncta number was quantified with Fiji software. Peroxisome distribution was analyzed by Icy software. Deconvolution of STED images was performed using Huygens software(Scientific Volume Imaging) with the Huygens classical maximum likelihood estimation deconvolution algorithm.

**Statistical analyses**

All statistical analyses, except hormone secretion, lifespan, and starvation survival-related experiments, were performed using Student’s *t*-test, and p<0.05 was considered a significant difference. For the hormone secretion test assay, The Chi-square test was utilized to calculate the p-value and p<0.05 was considered a significant difference. For lifespan and starvation survival-related assays, log-rank (mantel-Cox) tests were used to calculate the *p* values, and p<0.05 was considered a significant difference.
